# NON-REDUNDANT ROLES OF COPPER TRANSPORTERS ATP7A AND ATP7B IN NORADRENERGIC SIGNALING

**DOI:** 10.64898/2026.01.15.699802

**Authors:** Shubhrajit Roy, Yu Wang, Susan Aja, Niharika Sinha, Andrew Crawford, Keith MacRenaris, Alexa Wade, Kyungdo Kim, Abigael Muchenditsi, Lauryn Torres-Hernandez, Chan Hyun Na, Anastasia Kralli, Michael Petris, Anna Taylor, Thomas O’Halloran, Cay Tressler, Kristine Glunde, Svetlana Lutsenko

## Abstract

Menkes disease and Wilson disease are debilitating neurometabolic disorders caused by mutations in the copper (Cu) transporters ATP7A and ATP7B, respectively. In either disease, normalization of systemic Cu levels often does not eliminate neurological deficits, suggesting dysregulated Cu homeostasis within vulnerable neuronal populations. However, the specific roles of ATP7A and ATP7B and the extent of their functional redundancy in neurons remain poorly defined. Here, we selectively deleted Atp7a or Atp7b in noradrenergic neurons, which express both transporters and require Cu for catecholamine biosynthesis. ATP7A deletion reduced Cu levels in the locus coeruleus, disrupted dopamine-β-hydroxylase localization, impaired norepinephrine synthesis, and induced proteomic signatures of defective vesicular trafficking and proteostasis, culminating in neurodegeneration and impaired regulation of energy balance and adaptive thermogenesis. In contrast, ATP7B deletion preserved Cu levels but altered intracellular Cu utilization, resulting in catecholamine imbalance, α-synuclein upregulation, aberrant dopamine-β-hydroxylase distribution, and dysregulated thermogenesis. These findings establish ATP7A and ATP7B as non-redundant regulators of noradrenergic function within neural circuits governing metabolic and energy homeostasis and provide a mechanistic framework for persistent neurological pathology independent of systemic Cu levels.

## INTRODUCTION

Copper (Cu) is essential for human development and health. By serving as an enzymatic cofactor, Cu supports mitochondrial respiration, neurotransmitter synthesis, antioxidant defense, and other cellular processes, making adequate Cu availability especially critical for the nervous system. Copper-transporting ATPases ATP7A and ATP7B are major regulators of Cu homeostasis in cells and tissues: they deliver Cu to the cuproenzymes within the secretory pathway and export excess Cu to maintain intracellular Cu balance(Lutsenko et al., 2024). Loss-of-function mutations in ATP7A cause Menkes disease (MD), a fatal neurodegenerative disorder characterized by systemic Cu deficiency, widespread inactivation of essential cuproenzymes, catecholamine misbalance, and profound developmental delays(Fujisawa et al., 2022; Ramani and Parayil Sankaran, 2024; Tumer and Moller, 2010). Mutations in *ATP7B* cause Wilson disease (WD), in which Cu accumulates in the liver, brain, and other tissues, resulting in hepatic failure, movement disorders, psychiatric symptoms, and metabolic disturbances, including changes in catecholamines (Czlonkowska et al., 2018).

MD and WD patients have neurological deficits that are traditionally attributed to altered Cu levels in the brain, i.e. low Cu in MD and elevated Cu in WD (Dusek et al., 2019; Fujisawa et al., 2022). Indeed, treatments aimed at restoring Cu levels can improve neuronal function in either disease (Mohr and Weiss, 2019; Sarkar et al., 1993). However, despite early Cu supplementation in MD, neurological improvement remains incomplete. Similarly, Cu chelation in WD corrects but does not fully reverse neurologic symptoms and can even trigger worsening (Litwin et al., 2024). These results indicate that mechanisms beyond systemic Cu imbalance contribute to neuropathology. How specific neurons regulate their Cu homeostasis and what happens when these homeostatic mechanisms are disrupted remain poorly understood.

To uncouple the contribution of ATP7A and ATP7B to neuronal physiology at the cellular and circuit levels, we focused on noradrenergic neurons. These neurons express both ATP7A and ATP7B ((Lutsenko et al., 2019; Schmidt et al., 2018; Washington-Hughes et al., 2023) and **Fig. 1a**), rely on Cu-dependent dopamine-β-hydroxylase (DBH) for biosynthesis of norepinephrine (NE) (their primary effector molecule), and control organism-level processes such as adaptive thermogenesis. Adaptive thermogenesis is mediated by brown adipose tissue (BAT) and inducible beige adipocytes within white adipose tissue (WAT). BAT is the major thermogenic organ, whereas beige adipocytes expand thermogenic capacity in response to environmental stimuli (2–5). NE release from noradrenergic neurons activates the β-adrenergic receptors on adipocytes, triggering lipolysis and inducing uncoupling protein-1 (UCP1), which dissipates the mitochondrial proton gradient to generate heat. Cu is required for this pathway as a cofactor for cytochrome c oxidase, which establishes the mitochondrial proton gradient, and for DBH, which converts dopamine to NE. Thus, systemic Cu imbalance (either deficiency or excess) simultaneously perturbs neuronal signaling, mitochondrial bioenergetics, and peripheral metabolism, making it difficult to discern the specific neuronal functions of ATP7A and ATP7B.

**Figure 1.**
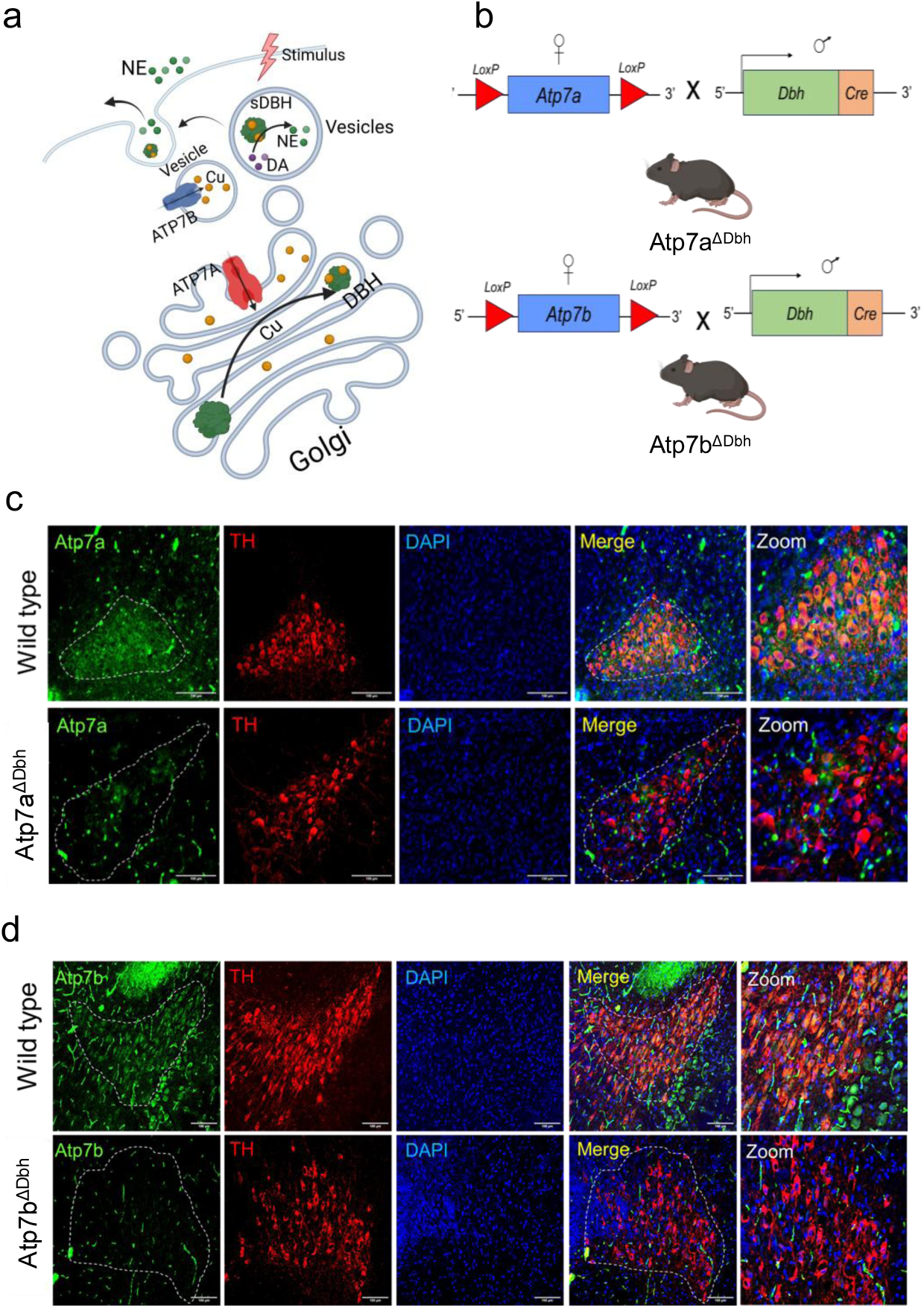
Targeted inactivation of *Atp7a* and *Atp7b* in noradrenergic neurons. (**a**)The schematic of predicted functions of the ATP-driven Cu transporters in noradrenergic neurons. ATP7A is located in the trans-Golgi network and delivers Cu cofactor to DBH; ATP7B is in cytoplasmic vesicles, where it sequesters Cu from the cytosol. Cu-bound DBH is located in secretory granules where it converts dopamine (DA) to norepinephrine (NE), which is then released upon neuronal activation. (**b**) Strategy for generating *Atp7a^ΔDbh^* and *Atp7b^ΔDbh^* mice: *Atp7a^LoxP/LoxP^* or *Atp7b^LoxP/LoxP^* female mice were crossed with male mice expressing Cre-recombinase under the DBH promoter (*DBH*-Cre mice) for targeted deletion in noradrenergic neurons. (**c,d)** Coronal brain sections from wild-type control and *Atp7a^ΔDbh^* **(c)** or *Atp7b^ΔDbh^* mice **(d)** were co-immunostained for tyrosine hydroxylase (Th) (Red), a marker of noradrenergic neurons, and either Atp7a (Green) or Atp7b (Green). In *Atp7a^ΔDbh^* and *Atp7b^ΔDbh^* mice, the respective transporter signal was undetectable in Th-positive neurons (outlined by dotted line), indicating successful deletion within the noradrenergic cell population. 3 coronal brain sections for n=3; animals were analyzed per genotype.

In this study, we clarified these functions by individually deleting Atp7a or Atp7b in noradrenergic neurons in mice and characterizing biochemical, cellular, and physiological consequences of their deletion. We found that Atp7a activity is essential for neuronal survival, noradrenergic signaling, and adaptive thermogenesis, independently of systemic Cu status. These findings reveal a neuronal contribution to thermogenic failure in MD and establish a role for neuronal Cu homeostasis in maintaining energy balance. We also uncovered a previously unrecognized role for Atp7b in supporting thermogenic responses and demonstrated dependence of DBH abundance on cellular Cu levels, especially at the cell periphery. Beyond their relevance to MD and WD, our results have implications for mechanistic studies of other neurodegenerative disorders for which Cu imbalance has been reported, such as Fatal Congenital Copper Deficit, Parkinson’s disease, Alzheimer’s disease, and aging.

## Results

### Targeted inactivation of Atp7a but not Atp7b alters Cu levels in the locus coeruleus (LC)

To investigate the physiological importance of Cu homeostasis in noradrenergic (NE) neurons, we individually deleted *Atp7a* or *Atp7b* in Dbh-expressing cells by crossing *Atp7a^LoxP/LoxP^* or *Atp7b^LoxP/LoxP^* females, respectively, with transgenic males expressing *Cre* recombinase under the *Dbh* promoter (Yamaguchi et al., 2018) (**Fig.1b**). Offspring were genotyped by PCR to verify the presence of the *Cre* allele and confirm successful recombination (**Fig. S1**). Both knockouts (*Atp7a^ΔDbh^* and *Atp7b^ΔDbh^*) were born at expected Mendelian frequencies; neither showed postnatal lethality or the lifespan reduction when compared to their respective *LoxP/LoxP* controls, indicating that the loss of either Cu transporter in NE neurons does not compromise animal survival under basal conditions.

Locus coeruleus (LC) is the brain’s main source of norepinephrine (Schwarz and Luo, 2015) and an anatomically distinct and easily identifiable cluster of NE neurons. Immunostaining of coronal brain sections containing LC confirmed deletion of Atp7a or Atp7b proteins specifically in the NE neurons (**Fig. 1c-d**). The deletion of either Atp7a or Atp7b did not significantly alter the total Cu content of the brain, as determined by atomic absorption spectroscopy of whole brain homogenates (**Fig. 2a**). Similarly, the peripheral tissues, such as liver and adipose tissue, had Cu levels similar to those of respective *LoxP/LoxP* controls in both knockout strains (**Fig. 2b**). Thus, systemic Cu balance is unaffected by the deletion of either Atp7a or Atp7b in Dbh-expressing cells, allowing us to examine specific consequences of Cu dis-homeostasis in neurons.

**Figure 2:**
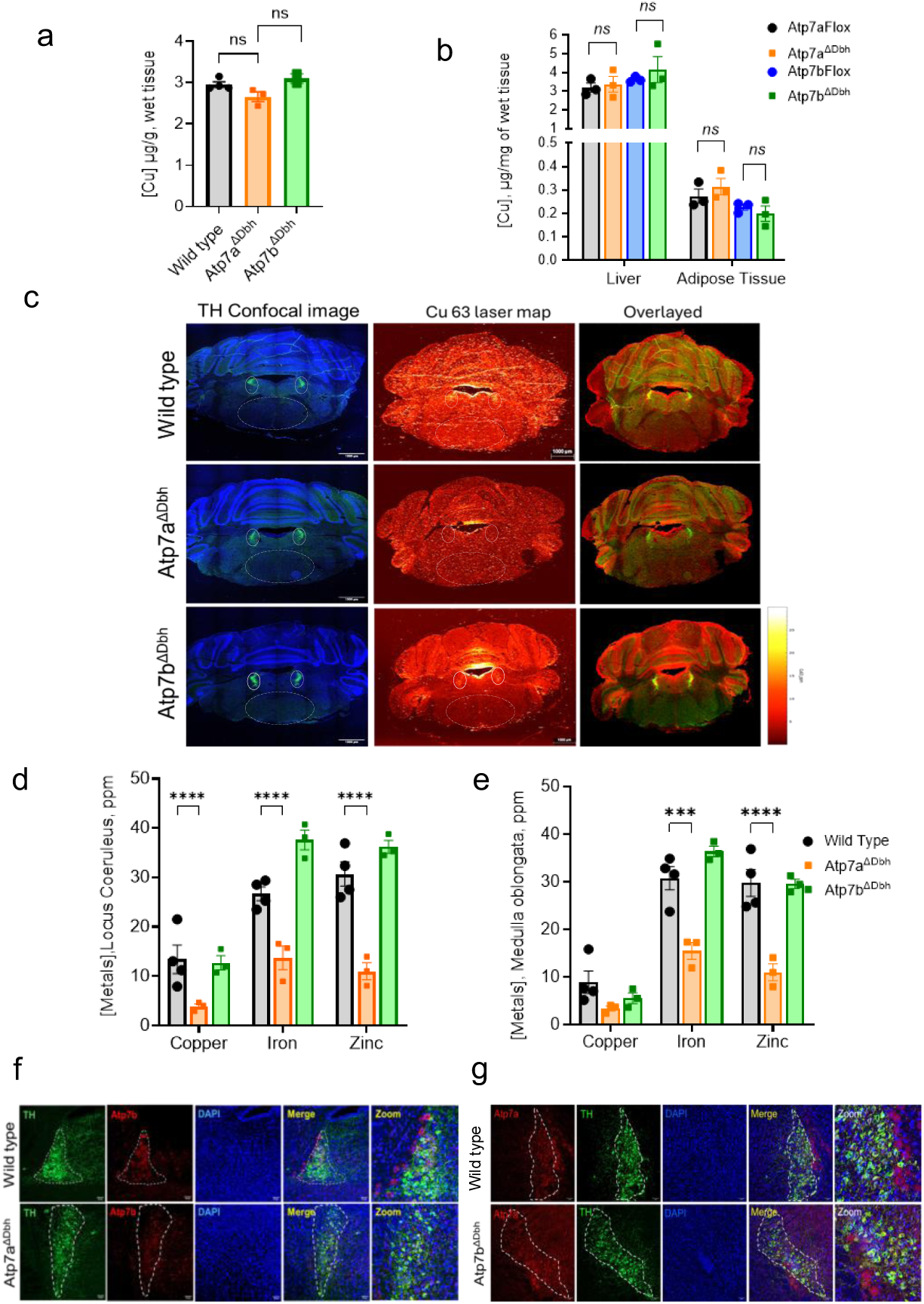
The effects of *Atp7a* and *Atp7b* deletion on Cu levels in locus coeruleus. **(a)** Analysis of Cu content in whole brain homogenates by atomic absorption spectroscopy found no difference for the age-matched 20-week-old wild-type, *Atp7a^ΔDbh^*, and *Atp7b^ΔDbh^* mice **(b)** Cu levels in peripheral tissues (liver, adipose) are unchanged across the genotypes. (**c**) Representative coronal brain sections analyzed by laser ablation inductively coupled plasma mass spectrometry (LA-ICP-MS) are shown, with metal distribution maps overlaid on tyrosine hydroxylase immunofluorescence confocal images (TH is green, nuclei are blue). Less intense Cu signal is apparent in the Th-positive region. (**d**) Quantitation of LA-ICP-MS signals in coronal brain sections reveals reduced Cu, Fe, and Zn levels in the locus coeruleus of Atp7a^ΔDbh^ mice compared to wild-type controls. (**e**) Metal levels in the medulla oblongata (ROI) (indicated by white dotted lines) are compared across wild-type, *Atp7a^ΔDbh^*, and *Atp7b^ΔDbh^* mice. (**f**) Immunostaining for Atp7b (red) in noradrenergic LC neurons (TH, green; LC outlined by the white dotted line) shows reduced Atp7b signal in *Atp7a^ΔDbh^* mice compared to wild type (Supplementary figure S2) (**g**) Immunostaining for Atp7a (red) in noradrenergic LC neurons (TH, green; LC outlined by white dotted line) shows no change in *Atp7b^ΔDbh^* mice compared to wild type. In all experiments, 3-4 mice were used for each genotype, ns = not significant; *p < 0.05; **p < 0.01; ***p < 0.001; ****p < 0.0001.

The NE neurons constitute only a small fraction of the total brain cell population, and a whole-brain analysis may not detect changes in their Cu content. Consequently, we examined Cu levels specifically within the LC region using laser ablation–inductively coupled plasma-time of flight-mass spectrometry (LA-ICP-TOF-MS). Adjacent sections were used for immunostaining of tyrosine hydroxylase (Th) to identify LC; the Th immunofluorescence images were then co-registered with the metal maps (**Fig. 2c and Fig.S2**). This analysis revealed a significant reduction of Cu levels in the LC of *Atp7a^ΔDbh^* mice relative to the age-matched 20-week-old wild-type controls (**Fig. 2d** and **Table S1**). The Cu levels outside of LC (such as *medulla oblongata*) also showed a mild decrease (**Fig. 2e**), indicating that the loss of Cu in *Atp7a^ΔDbh^* LC impacts the surrounding medullary region. Broader perturbations in metal homeostasis in the *Atp7a^ΔDbh^* LC were further highlighted by unexpected decreases in iron (Fe) and zinc (Karczewski et al.) (**Fig. 2d and 2e**). In contrast, neither *Atp7b^ΔDbh^*LC nor the surrounding areas showed significant changes in copper or other metals (**Fig. 2d and 2e**).

### Dbh protein abundance is reduced in Atp7a^ΔDbh^ and Atp7b^ΔDbh^ mouse brains

Since the biochemical functions of Atp7a and Atp7b are very similar, we investigated whether the abundance of either transporter changed to compensate for the loss of the other. Immunostaining of Atp7b in *Atp7a^ΔDbh^* LC reveals no increase in the Atp7b signal; in fact, the intensity of Atp7b staining appeared lower in *Atp7a^ΔDbh^* cells (**Fig. S3**). Similarly, Atp7a levels were not noticeably increased in *Atp7b^ΔDbh^* mice (**Fig. 2f-g**). By contrast, the abundance of Dbh protein changed significantly. The 20-week-old *Atp7a^ΔDbh^* whole^−^brain homogenates and LC both showed marked loss of the Dbh signal compared to control (**Fig. 3 a-d**). In younger *Atp7a^ΔDbh^* animals (4 weeks after birth), Dbh expression was also diminished (**Fig. 3e-f**), suggesting that low Dbh abundance is likely a direct result of Cu misbalance during brain development, rather than a secondary long-term change.

**Figure 3.**
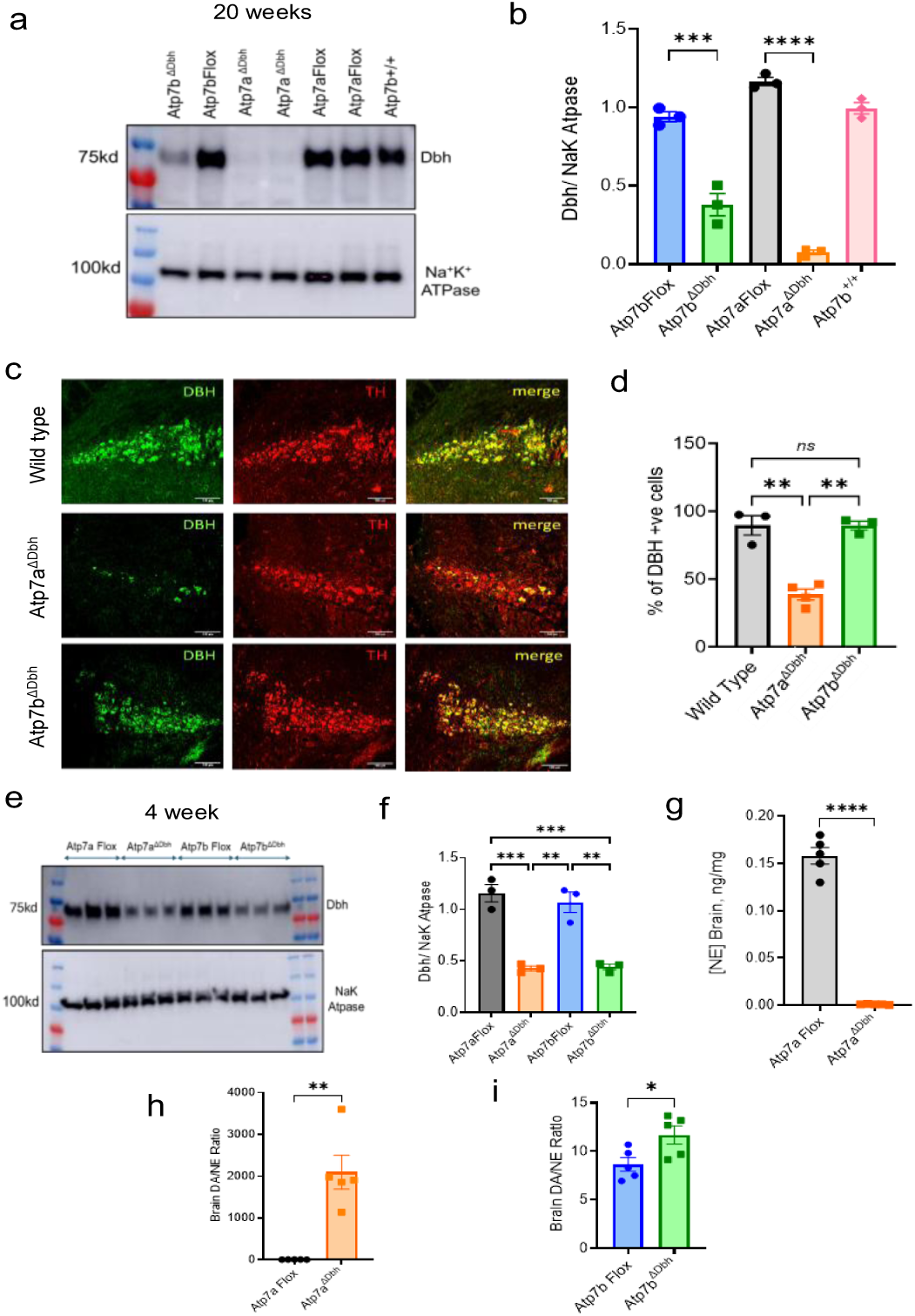
Inactivation of *Atp7a* or *Atp7b* in NE neurons reduces Dbh expression and alters catecholamine metabolism. (**a**) Representative immunoblots of the whole-brain homogenates show markedly reduced Dbh expression in 20-week-old *Atp7a^ΔDbh^* compared to *Atp7a^LoxP/LoxP^* (*Atp7aFlox*) and wild-type controls. Additional replicates are shown in the Supplementary figure S11. (**b**) Densitometry of DBH signal normalized to Na^+^/K^+^ ATPase confirms significantly reduced Dbh in 20-week-old *Atp7a^ΔDbh^* brains. (**c**) Representative coronal brain sections (10 μm) from 20-week-old mice immunostained for Dbh (green) and tyrosine hydroxylase (Th, red) highlight noradrenergic neurons in the locus coeruleus (LC); *Atp7a^ΔDbh^*brains display marked loss of Dbh signal compared to controls. (**d**) Dbh-positive neurons were normalized to the total number of TH-stained neurons in the LC. Three coronal brain sections from n=3 animals per genotype were analyzed. (**e**) Immunoblots at 4 weeks similarly show reduced Dbh expression in both *Atp7a^ΔDbh^* and *Atp7b^ΔDbh^*. (**f**) Densitometry of the Dbh signal normalized to Na^+^/K^+^ ATPase confirms significantly reduced Dbh in 4-week-old *Atp7a^ΔDbh^* and *Atp7b^ΔDbh^* brains. LC-MS metabolite analysis of whole-brain homogenates reveals a (**g**) decreased norepinephrine (NE) and (**h**) increased dopamine (DA)/NE ratio in *Atp7a^ΔDbh^* mice relative to *Atp7aFlox* controls (20 weeks, n=5). (**i**) increased DA/NE ratio in *Atp7b^ΔDbh^* brain homogenates relative to *Atp7bFlox* controls (20 weeks, n = 3). Data are mean ± SEM. Statistical analysis: unpaired two-tailed Student’s t-test with Welch’s correction; *p < 0.05, **p < 0.01. Data points for *Atp7aFlox, Atp7bFlox, Atp7a^ΔDbh^,* and *Atp7b^ΔDbh^* are shown in black, blue, orange, and green, respectively.

Unlike the *Atp7a^ΔDbh^* LC, the *Atp7b^ΔDbh^*brain sections that include LC showed no difference in Dbh staining compared to the control. Since the Dbh protein is produced in cell bodies and then transported to the cell periphery to synthesize NE in secretory granules, we also measured Dbh levels in the entire brain. In whole-brain homogenates from *Atp7b^ΔDbh^* animals, Dbh levels were significantly reduced - by about 50% - compared to the age-matched controls (**Fig. 3a-b** and **Fig.S4**). In both *Atp7a^ΔDbh^*and *Atp7b^ΔDbh^* mice, the reduction in Dbh levels was greater in whole brain homogenates than in the LC cell bodies, suggesting that Cu imbalance negatively impacts DBH delivery to cell periphery, likely diminishing NE synthesis/neurotransmission. Direct measurements of dopamine (DA) and norepinephrine (NE) confirmed a marked increase in the DA/NE ratio in *Atp7a^ΔDbh^* brains (**Fig. 3g-h**), indicative of impaired DA-to-NE conversion. A less dramatic, but statistically significant increase in the DA/NE ratio was observed in *Atp7b^ΔDbh^*brains (**Fig. 3g**). Serotonin levels remained unchanged in both *Atp7a^ΔDbh^* and *Atp7b^ΔDbh^* animals (**Fig. S5)**, indicating that Cu imbalance in the NE neurons specifically dysregulates NE/DA homeostasis without broadly affecting monoamine metabolism.

### Inactivation of Atp7a dysregulates cell adhesion and vesicular trafficking, and increases autophagy markers

To better understand the molecular effects of *Atp7a* and *Atp7b* deletions and compare these effects, we analyzed the LC proteomes of knockout animals and respective *LoxP/LoxP* controls (3 animals per group for each of four genotypes). The LC regions, identified by immunostaining for tyrosine hydroxylase (Th), were isolated using laser-capture microdissection and subjected to tandem mass tag (TMT)-labelling and liquid chromatography-mass spectrometry (**Fig.S6**). A total of 3,963 proteins were detected in all 12 samples, and their relative abundances were quantified. The LC with inactivated Atp7a showed a greater number of significantly changed proteins, 131 (1.2-fold change, q < 0.05), compared to the respective control, whereas *Atp7b^ΔDbh^* LC had 43 proteins changed (1.2-fold change, q < 0.05) (**Fig. 4a-d and Table S2**).

**Figure 4.**
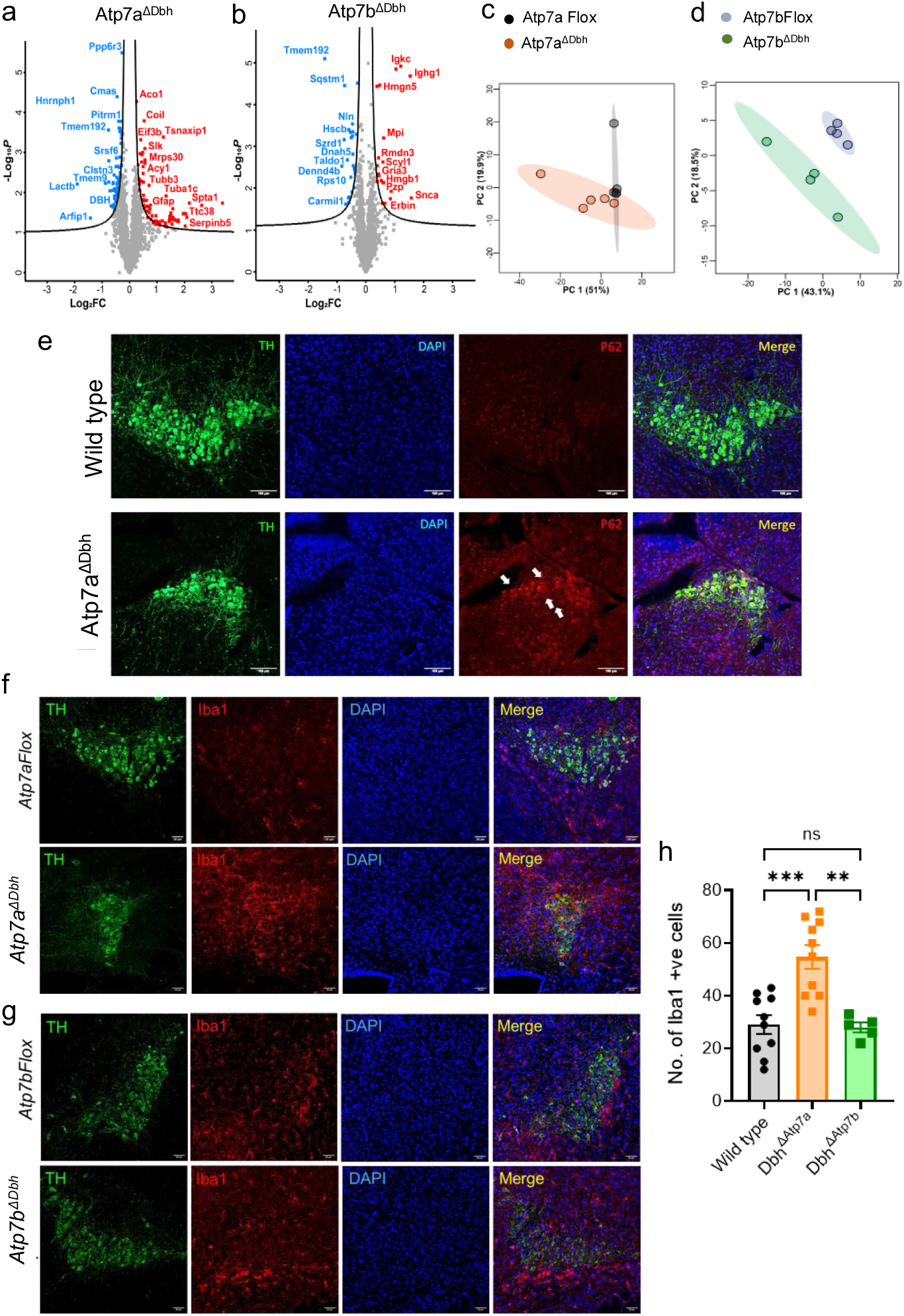
Loss of *Atp7a* but not *Atp7b* induces autophagic and neuroinflammatory responses in locus coeruleus. (a, b) Volcano plots showing differentially expressed proteins in *Atp7a^ΔDbh^* (**a**) and *Atp7b^ΔDbh^* (**b**) laser-dissected locus coeruleus compared to the respective *LoxP/LoxP* controls. (c, d) Principal component analysis of proteomic datasets demonstrates clear segregation between mutant and control samples for *Atp7a^ΔDbh^* (**c**) and *Atp7b^ΔDbh^* (**d**). (**e**) Immunofluorescence staining of coronal brain sections demonstrates accumulation of the autophagy adaptor p62 (red) in *Atp7a^ΔDbh^* neurons, colocalizing with tyrosine hydroxylase (TH, green). (f) Enhanced microglial activation in *Atp7a^ΔDbh^* LC, indicated by increased Iba1 immunoreactivity (red). (**g**) Iba1 staining in *Atp7b^ΔDbh^* LC is similar to controls. (**h**) Quantification of Iba1+ cells in the LC (n = 3 mice per genotype). Data are mean ± SEM. Statistical analysis: unpaired two-tailed Student’s t-test with Welch’s correction; *p < 0.05, **p < 0.01. Data points for *Atp7aFlox*, *Atp7bFlox*, *Atp7a^ΔDbh^,* and *Atp7b^ΔDbh^* are shown in black, blue, orange, and green, respectively.

In *Atp7a^ΔDbh^* LC, the most down-regulated proteins included tyrosine hydroxylase, Dbh, and catecholamine transporter Slc18a2 – all required for norepinephrine biosynthesis. We also observed a decrease in proteins involved in RNA synthesis (Rps24, Polr2m, Sf3b6) and lipid and energy balance (Slc25a3, Lactb), suggesting metabolic deficits in the neurons. The largest category of dysregulated proteins included proteins involved in cell adhesion, vesicular trafficking, neuronal guidance, and synapse formation (**Table 1 and Table S2**). Significant changes were also observed in proteins involved in inflammatory response (Serpinb1a, Nrlx1, Map3k7) and apoptosis/protein degradation (Slk, Tmem9, Tmem192, Ube2a, Ddrgk1). The list of all significantly changed proteins is included in **Table S2**. Together, these changes suggested a potential axonal atrophy and/or loss of neurons. To test this, we co-immuno-stained the LC-containing brain sections with antibodies against p62 (SQSTM1), which promotes the recruitment of ubiquitinated proteins to autophagosomes and tyrosine hydroxylase (Th) Although in the entire LC proteome, SQSTM1 was not significantly changed, in Th-positive neurons of *Atp7a^ΔDbh^* mice, SQSTM1/p62 signal was strongly increased, consistent with cellular stress and autophagy (**Fig. 4e**). Strong increase in the immunostaining of Iba, a marker of activated microglia/macrophages (**Fig. 4f-h**) further pointed to neuronal degeneration and an increased inflammatory response.

**Table 1.** The most significantly changed proteins (fold change >1.5, p-value<0.05) in *Atp7a*^Δ*DBH*^ locus coeruleus (proteins directly involved in NE metabolism are in blue)

| Symbol | Protein name | Fold Change | Function |
| --- | --- | --- | --- |
| **Upregulated in *Atp7a*<sup>ΔDBH</sup>** |  |  |  |
| VAC14 | VAC14 component of PIKFYVE complex | 2.49 | synthesis and turnover of PI(3,5)P2, endosomal trafficking |
| TSNAXIP1 | translin associated factor X interacting protein 1 | 2.47 | sperm motility and cilia function |
| NDEL1 | nudE neurodevelopment protein 1 like 1 | 1.76 | neuronal migration, mitosis, and the maintenance of cell structures |
| FRMD4A | FERM domain containing 4A | 1.74 | scaffolding, cell polarity |
| MEMO1 | mediator of cell motility 1 | 1.66 | cell migration |
| SCYL1 | SCY1 like pseudokinase 1 | 1.54 | regulating Golgi morphology and COPI trafficking |
| SCN2B | sodium voltage-gated channel beta subunit 2 | 1.5 | ion transport |
| **Downregulated in *Atp7a*<sup>ΔDBH</sup>** |  |  |  |
| CLSTN3 | calsyntenin 3 | 0.63 | postsynaptic adhesion molecule |
| ICA1 | islet cell autoantigen 1 | 0.63 | role in neurotransmitter secretion |
| POLR2M | RNA polymerase II subunit M | 0.62 | translation |
| C12orf43 | chromosome 12 open reading frame 43 | 0.62 | regulation of Wnt signaling pathway |
| CKAP4 | cytoskeleton associated protein 4 | 0.61 | cell adhesion and migration |
| DBH | dopamine beta-hydroxylase | 0.61 | NE synthesis |
| CARMIL1 | capping protein regulator and myosin 1 linker 1 | 0.6 | role in cells protrusions |
| AASDHPPT | amino adipate-semialdehyde dehydrogenase-phosphopantetheinyl transferase | 0.58 | biosynthesis of lysine and fatty acids. |
| TMEM192 | transmembrane protein 192 | 0.56 | lysophagy |
| SLC25A31 | solute carrier family 25 member 31 | 0.56 | mitochondria function, energetics |
| ARFIP1 | ARF interacting protein 1 | 0.55 | biogenesis of secretory granules at the trans-Golgi network |
| TH | tyrosine hydroxylase | 0.53 | DOPA synthesis |
| SLC18A2 | solute carrier family 18 member A2 | 0.52 | monoamine transporter |
| LACTB | lactamase beta | 0.37 | mitochondria lipid metabolism |

**Table 2.** The most significantly changed proteins (fold change >1.5, p-value<0.05) in *Atp7b*^Δ*DBH*^ locus coeruleus.

| Symbol | Protein name | Fold Change | Function |
| --- | --- | --- | --- |
| **Upregulated in *Atp7b*<sup>ΔDBH</sup>** |  |  |  |
| SNCA | synuclein alpha | 2.9896985 | regulation of synaptic functions |
| IgHg | Immunoglobulin heavy chain (gamma polypeptide) | 2.91399823 | long term immunity |
| IGKC | immunoglobulin kappa constant | 2.05765342 | antibody stabilization |
| ALB | albumin | 1.83400809 | inhibition of apoptosis |
| ERBIN | erbB2 interacting protein | 1.80875876 | cell polarity, apoptosis |
| SERPIN1A | serine peptidase inhibitor, clade A, member 1A | 1.59107297 | protective, anti-inflammatory tole |
| MVD | mevalonate diphosphate decarboxylase | 1.5822746 | lipid synthesis, synaptic plasticity |
| APOA1 | apolipoprotein A1 | 1.54114222 | cholesterol transport |
| MPI | mannose phosphate isomerase | 1.53900722 | glycosylation |
| Pzp | PZP, alpha-2-macroglobulin like | 1.53368266 | inhibitor of NGF-dependent neuronal outgrowth |
| SCYL1 | SCY1 like pseudokinase 1 | 1.50316148 | regulating Golgi morphology and COPI trafficking |
| **Downregulated in *Atp7b*<sup>ΔDBH</sup>** |  |  |  |
| TALDO1 | transaldolase 1 | 0.63683874 | pentose phosphate pathway |
| CARMIL1 | capping protein regulator and myosin 1 linker 1 | 0.6172813 | actin polymerization |
| SQSTM1 | sequestosome 1 | 0.59832448 | autophagy receptor |
| SZRD1 | SUZ RNA binding domain 1 | 0.59295725 | nonsense-mediated decay |
| DENND4B | DENN domain containing 4B | 0.55980644 | Rab10 activator, vesicle trafficking |
| TMEM192 | transmembrane protein 192 | 0.37241937 | lysophagy |

### Inactivation of Atp7b affects the LC proteome differently from Atp7a deletion

Changes in the proteome of *Atp7b^ΔDbh^* LC were markedly different. Not only were fewer proteins changed, but an overlap between the differentially expressed proteins in *Atp7a^ΔDbh^* and *Atp7b^ΔDbh^*LC was minimal. Commonly changed proteins include TMEM192 (lysosomal function)(Park et al., 2025), which was down-regulated in both strains, and Scyl1(Kaeser-Pebernard et al., 2022) (involved in Golgi maintenance and vesicular trafficking), which was commonly upregulated. The abundances of Dbh and Th were not changed in Atp7b KO in agreement with the LC immunostaining data. Also, upregulation of Iba1 levels was not observed (**Fig. 4g-h**). Instead, we found a highly significant increase in the abundance of α-synuclein (3-fold, *p-value* = 0.0017), which was confirmed by immunostaining (**Fig. S7)**, pointing to potential changes in neurotransmission in NE neurons lacking Atp7b. Changes in the abundances of Scyl1, Dennd4b, Rmdn3, and Mylk further pointed to altered vesicle trafficking and recycling as well as dysregulation of axonal outgrowth (Burman et al., 2008). Other categories of proteins with changed abundance included metabolic enzymes and proteins involved in immune response. Sqstm1 (Gallagher and Holzbaur, 2023; Tominaga et al., 2017) was downregulated, which coincides with the upregulation of Pzp, a protein that facilitates protein folding (Cater et al., 2019). Taken together, these data suggested that inactivation of Atp7b in NE neurons leads to metabolic adaptation and potentially altered secretory vesicular trafficking, but, unlike Atp7a, this deletion does not cause significant neuronal loss.

### Atp7a^ΔDbh^ and Atp7b^ΔDbh^ mice show distinct metabolic changes

To evaluate the functional consequences of noradrenergic Atp7a or Atp7b deletion, we first examined the known physiological effects of NE (Smythe et al., 1984; Walters et al., 1997). Previously, *Dbh^−/−^* mice that lack NE were shown to have hypoglycemia and hyperinsulinemia, implicating catecholaminergic circuits in metabolic control (Arnold et al., 2017). Consequently, we first examined glucose homeostasis in our knockouts. At steady-state, *Atp7a^ΔDbh^* mice showed hypoglycemia, whereas *Atp7b^ΔDbh^* did not (**Fig. 5a**). Following intraperitoneal glucose administration, *Atp7a^ΔDbh^* mice had lower blood glucose levels at all time points. Lower levels of blood glucose in *Atp7a^ΔDbh^* mice suggested either increased glucose clearance or impaired glucose production by the liver. To distinguish between these possibilities, we performed an insulin tolerance test and a pyruvate tolerance test, respectively. Although glucose levels at 30-, 60-, and 90 minutes post-insulin administration were lower in *Atp7a^ΔDbh^* animals, the kinetics of glucose clearance was unchanged (**Fig. 5b**). In contrast, following pyruvate injection, *Atp7a^ΔDbh^* mice showed less glucose at 15, 30, 60, and 90 minutes, compared to control, indicative of impaired hepatic glucose production.

**Figure 5:**
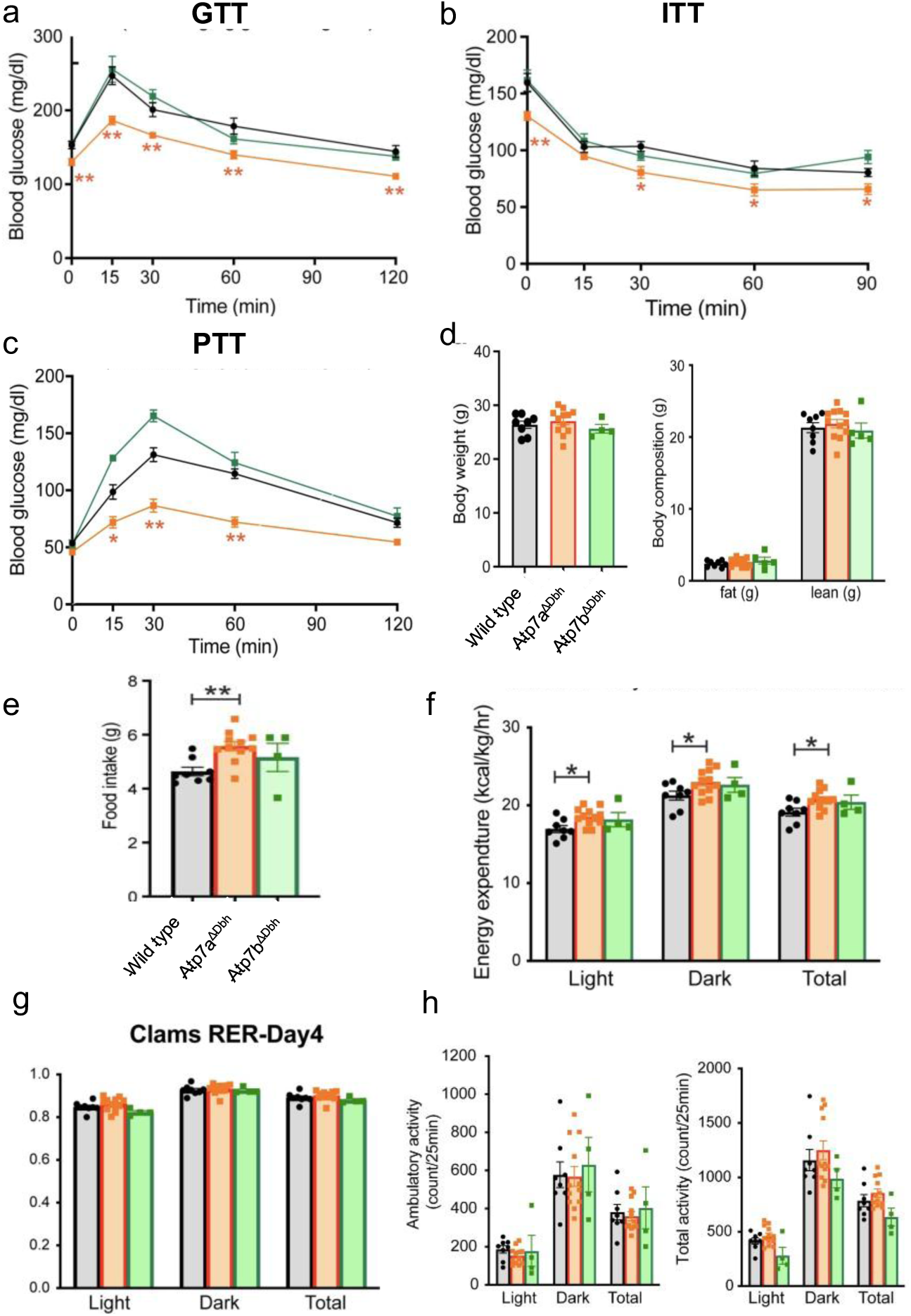
Altered glucose metabolism, enhanced energy expenditure, and food intake in *Atp7a^ΔDbh^* mice. (**a**) Glucose tolerance test (GTT) shows reduced blood glucose levels in *Atp7a^ΔDbh^* mice compared to the wild-type controls. (**b**) Insulin tolerance test (ITT) shows that *Atp7a^ΔDbh^* mice respond to insulin and remain hypoglycemic. (**c**) The pyruvate tolerance test (PTT) demonstrates impaired gluconeogenesis in *Atp7a^ΔDbh^* mice. (**d**) Body weight and composition (including fat and lean mass percentages) showed no significant differences between genotypes. (e) Daily food intake increased in *Atp7a^ΔDbh^* mice. (**f**) Indirect calorimetric analysis reveals that energy expenditure (normalized to lean mass or body weight) increases in *Atp7a^ΔDbh^* mice during light, dark, and whole-day (total) periods. (**g**) Respiratory exchange ratio (RER) shows no significant difference between the genotypes. (**h**) Ambulatory and total activity counts over light, dark, and whole-day periods show no significant differences between groups. Data are presented as mean ± SEM; p < 0.05; p < 0.01; p < 0.001; p < 0.0001; ns, not significant (n=5-12 mice per genotype)

*Atp7b^ΔDbh^* mice’s responses to glucose or insulin administration were similar to controls (**Fig. 5a**). In the pyruvate tolerance test, *Atp7b^ΔDbh^* animals initially showed a higher surge of blood glucose than controls, but their glucose levels returned to normal levels over time.

Despite metabolic changes, body weight, fat mass, and lean mass in *Atp7a^ΔDbh^* and *Atp7b^ΔDbh^* mice were similar to controls (**Fig. 5d**). Notably, *Atp7a^ΔDbh^* mice ate more food (hyperphagia) (**Fig. 5e**), along with a significantly increased oxygen consumption (VO₂) and carbon dioxide production (VCO₂), indicating a higher metabolic rate (**Fig. S8**). This rise in energy expenditure (**Fig. 5f**) occurred independently of changes in ambulatory or overall physical activity during both light and dark phases (**Fig. 5g, h**). Therefore, the Atp7a function in NE neurons is essential for NE signaling that regulates glucose balance and energy use. In contrast, *Atp7b^ΔDbh^* mice exhibited a modest reduction in locomotor activity without corresponding changes in metabolic rate (**Fig. 5h**); these differences were not statistically significant.

### Cu misbalance in NE neurons is associated with BAT metabolic quiescence

To identify physiological processes in *Atp7a^ΔDbh^* mice that require increased expenditure of metabolic energy, we examined their adaptive thermogenesis, which is an NE-dependent process (Garside et al., 2022). Under standard temperature conditions (∼22-24°C), both *Atp7a^ΔDbh^* and *Atp7b^ΔDbh^* mice displayed a warmth-seeking behavior, characterized by frequent and prolonged burrowing beneath nestlets within their cages (**Fig.6a**). This behavior was absent in the respective *LoxP/LoxP* controls, which remained active and exposed (**Fig. 6a**). The behavior to seek insulation suggested impaired thermoregulatory capacity or heightened cold sensitivity in *Atp7a^ΔDbh^* and *Atp7b^ΔDbh^* mice, even at ambient temperatures.

**Figure 6:**
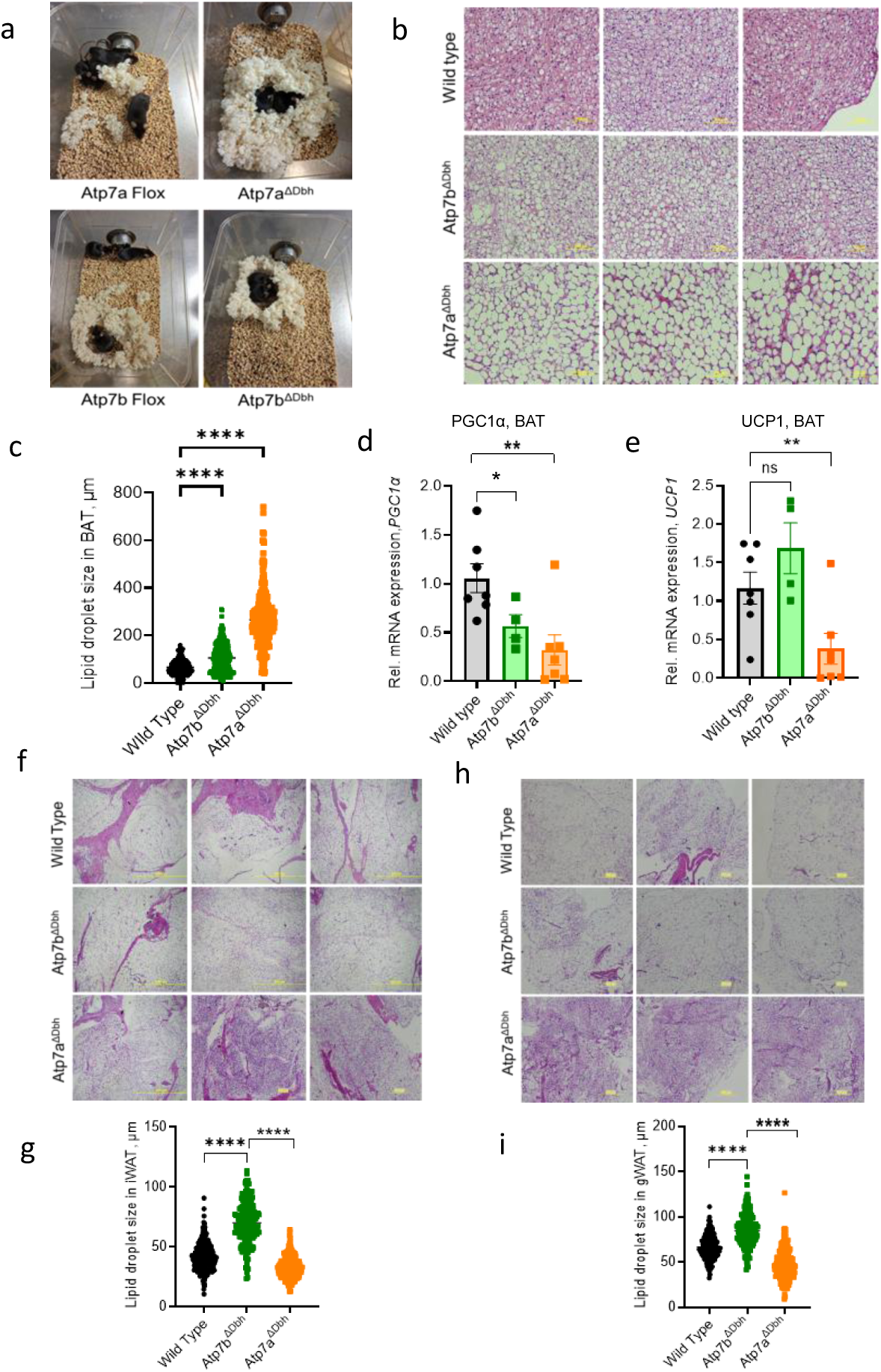
Altered non-shivering thermogenesis and adipose tissue morphology and thermogenic gene expression in *Atp7a^ΔDbh^* and *Atp7b^ΔDbh^*. **(a)** Images illustrating the warmth-seeking behaviour of *Atp7a^ΔDbh^* and *Atp7b^ΔDbh^* mice compared to the *Atp7aFlox* and *Atp7bFlox* animals. (**b**) Hematoxylin and eosin (H&E)-stained sections of brown adipose tissue (BAT) show enlarged adipocytes in *Atp7a^ΔDbh^* and *Atp7b^ΔDbh^* mice relative to the wild type. (c) Quantification of BAT adipocyte size confirms a significant increase in adipocyte size in both mutants. (d–e) Relative mRNA expression of thermogenesis-related genes in BAT: (**d**) *PGC1α* and (**e**) *UCP1* are reduced in *Atp7a^ΔDbh^* compared to wild type. (**f**) H&E-stained sections of inguinal white adipose tissue (iWAT) reveal enlarged adipocytes in *Atp7a^ΔDbh^*. (**g**) Quantification of iWAT adipocyte size demonstrates significantly increased adipocyte size in *Atp7a^ΔDbh^* mice compared to wild type. (h) H&E-stained sections of gonadal white adipose tissue (gWAT). (**i**) Quantification of gWAT adipocyte size shows significantly increased adipocyte size in *Atp7a^ΔDbh^* mice compared to wild type. Data represent mean ± SEM. Statistical analysis: unpaired Student’s t-test; ns = not significant; *p < 0.05, **p < 0.01, ****p < 0.0001.

To evaluate thermoregulatory capacity, we examined the major thermogenic tissue, BAT. We also analyzed WAT (including anatomically distinct inguinal (iWAT) and gonadal (gWAT) depots), which can adopt thermogenic characteristics under certain physiological stimuli (Finlin et al., 2018; Huang et al., 2017). Histological analysis of BAT from *Atp7a^ΔDbh^* mice revealed enlarged lipid droplets, suggesting impaired lipid utilization (**Fig. 6b**). A milder increase in lipid droplet size was also observed in *Atp7b^ΔDbh^* mice (**Fig. 6c**). The mRNA levels of thermogenic markers *PGC1α* and *UCP1* were significantly lower in BAT of *Atp7a^ΔDbh^* animals, consistent with the suppressed NE signaling to β-adrenergic receptors and decreased thermogenic capacity (**Fig. 6d&e**). In *Atp7b^ΔDbh^* BAT, the *PGC1α* mRNA levels were significantly lower than in control tissue, whereas the UCP1 transcript was unchanged, suggesting that some thermogenic capacity was preserved in these mice. In agreement with the histological findings, BAT from *Atp7a^ΔDbh^* mice weighed significantly more than BAT of wild-type controls, whereas *Atp7b^ΔDbh^* mice showed only a mild, non-significant increase in the BAT weight (**Fig. S9**).

Despite impaired lipid mobilization in BAT, the total body weight, fat mass, and lean mass of knockout mice were comparable to controls (**Fig. 5d**), prompting us to investigate whether WAT depots underwent compensatory remodeling (beiging). Inguinal and gonadal WAT from *Atp7a^ΔDbh^* mice showed multilocular, smaller lipid droplet size when compared to controls, resulting in darker eosinophilic cytoplasm in H&E staining. These histological findings suggested beiging of WAT, a morphological feature consistent with a compensatory thermogenic activation (**Fig. 6f-i**). A similar, although less robust phenotype was observed in *Atp7b^ΔDbh^* mice (**Fig. 6f-i**).

### Atp7a^ΔDbh^ and Atp7b^ΔDbh^ mice fail to maintain body temperature during acute cold exposure

To directly test how well *Atp7a^ΔDbh^*and *Atp7b^ΔDbh^* mice maintain their core body temperature, we first acclimated them to thermoneutral conditions for 10 days and then exposed mice to cold (6 °C). In response to cold, the core body temperature of both *Atp7a^ΔDbh^* animals and respective controls declines (**Fig. 7a**). After the initial drop, the control mice maintained their body temperature. In contrast, the body temperature of *Atp7a^ΔDbh^* mice dropped below the critical threshold (28°C) by 5 hours, which terminated the experiment.

**Figure 7:**
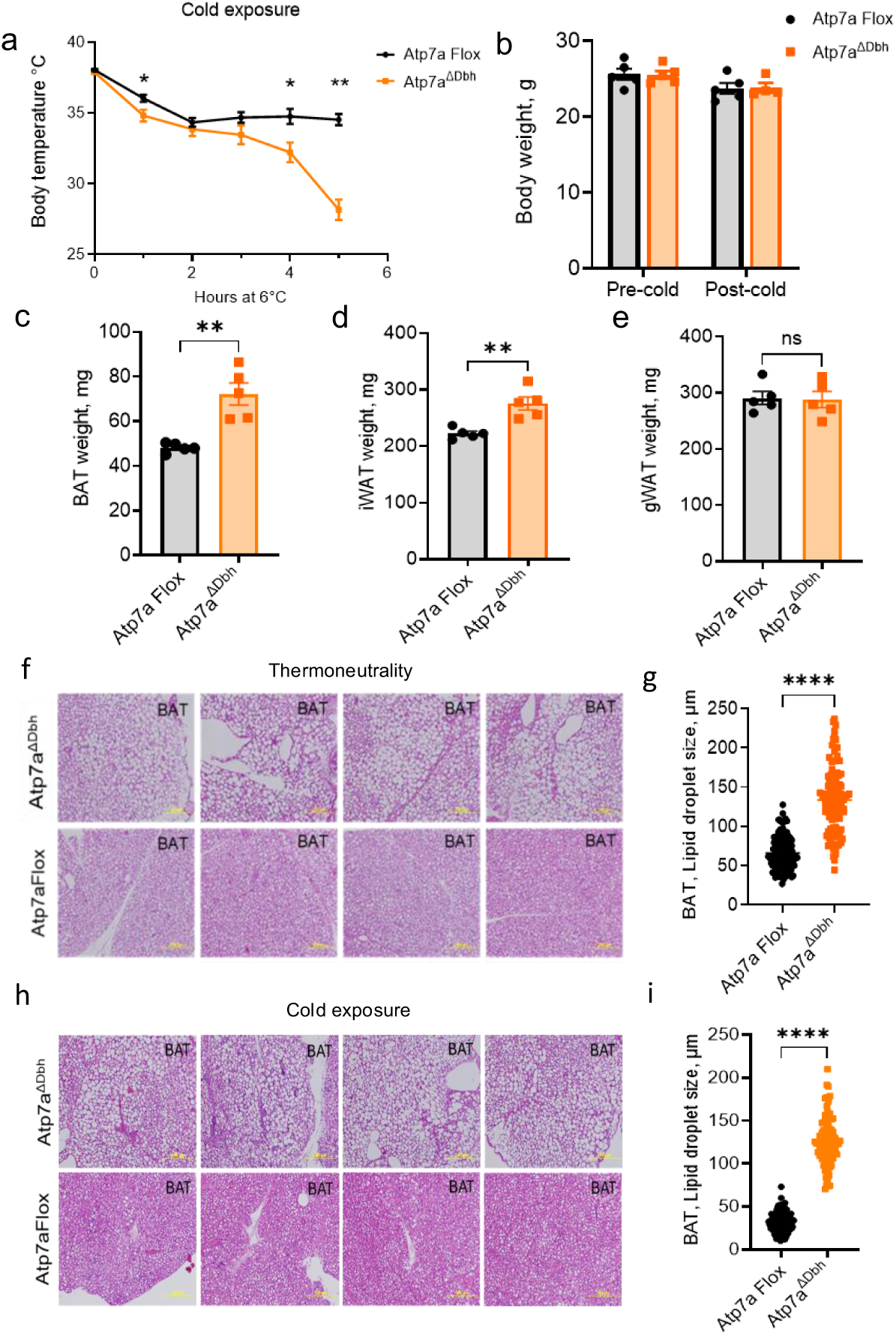
*Atp7a^ΔDbh^* animals exhibit severe hypothermia under acute cold stress. **(a)** Twelve-week-old *Atp7a^ΔDbh^* male mice and *Atp7aFlox* controls (n = 5-6) were exposed to acute cold stress (6°C) after prolonged thermoneutral housing (30°C for 10 days). Unlike the age-matched Atp7aFlox, *Atp7a^ΔDbh^* mice were unable to maintain core body temperature (>30°C). (**b**) The body weight of Atp7a^ΔDbh^ and Atp7aFlox controls showed no significant difference before and after cold exposure. (**c**) The weight of brown adipose tissue (BAT) in *Atp7a^ΔDbh^* animals is significantly higher compared to controls. (**d, e**) Gonadal white adipose tissue (gWAT) weight did not differ significantly between *Atp7a^ΔDbh^* and *Atp7aFlox* controls. (**f**) Representative hematoxylin and eosin (H&E) - stained sections of BAT from *Atp7aFlox* and *Atp7a^ΔDbh^* mice, maintained in thermoneutrality (30 °C for 10 days), reveal enlarged adipocytes indicative of defective lipolysis in mutant animals. (g) Quantification confirms significant increases in adipocyte size in BAT. (**h**) H&E-stained sections of BAT isolated from *Atp7a^ΔDbh^* and Atp7aFlox controls following acute cold stress at 6 °C for 5 hours show enlarged lipid droplets in BAT compared to the *Atp7bFlox* controls. (**i**) Quantification supports these observations. Data are presented as mean ± SEM. Statistical significance was determined by unpaired Student’s t-test: *p* < 0.05, p < 0.01, p < 0.0001; ns = not significant.

The body weights of control and *Atp7a^ΔDbh^* mice were similar either pre-cold or post-cold (**Fig. 7b**). However, the *Atp7a^ΔDbh^*BAT was significantly heavier after cold exposure than the BAT of *Atp7a^LoxP/LoxP^* controls, suggesting that *Atp7a^ΔDbh^*BAT could not utilize fat stores (**Fig. 7c** and **Fig. S10**). In support of impaired lipid mobilization, *Atp7a^ΔDbh^* BAT exhibited enlarged lipid droplets and defective lipolysis (**Fig. 7h–i**). BAT dysfunction in *Atp7a^ΔDbh^* mice was apparent even under prolonged thermoneutral conditions (30 °C, 10 days) (**Fig. 7f-g** and **Fig. S11**), pointing to a severe deficit in NE signaling.

In response to cold exposure, *Atp7b^ΔDbh^* animals also exhibited a significant decrease in core body temperature at 2h and 5h. However, they survived and at later time points were able to maintain a core body temperature comparable to that of the *Atp7b^LoxP/LoxP^* controls (**Fig. 8a**). Their weight at thermoneutrality trended to be lower than in controls, but the pre- and post-exposure body weights showed no significant differences (**Fig. 8b**). After cold exposure, *Atp7b^ΔDbh^* BAT weighed more than control BAT, whereas other adipose depots remained unchanged (**Fig. 8c-e** and **Fig. S10**). Taken together, these findings identify *Atp7a* as a key player in the noradrenergic regulation of adaptive thermogenesis, with *Atp7b* playing a more limited, likely fine-tuning role.

**Figure 8.**
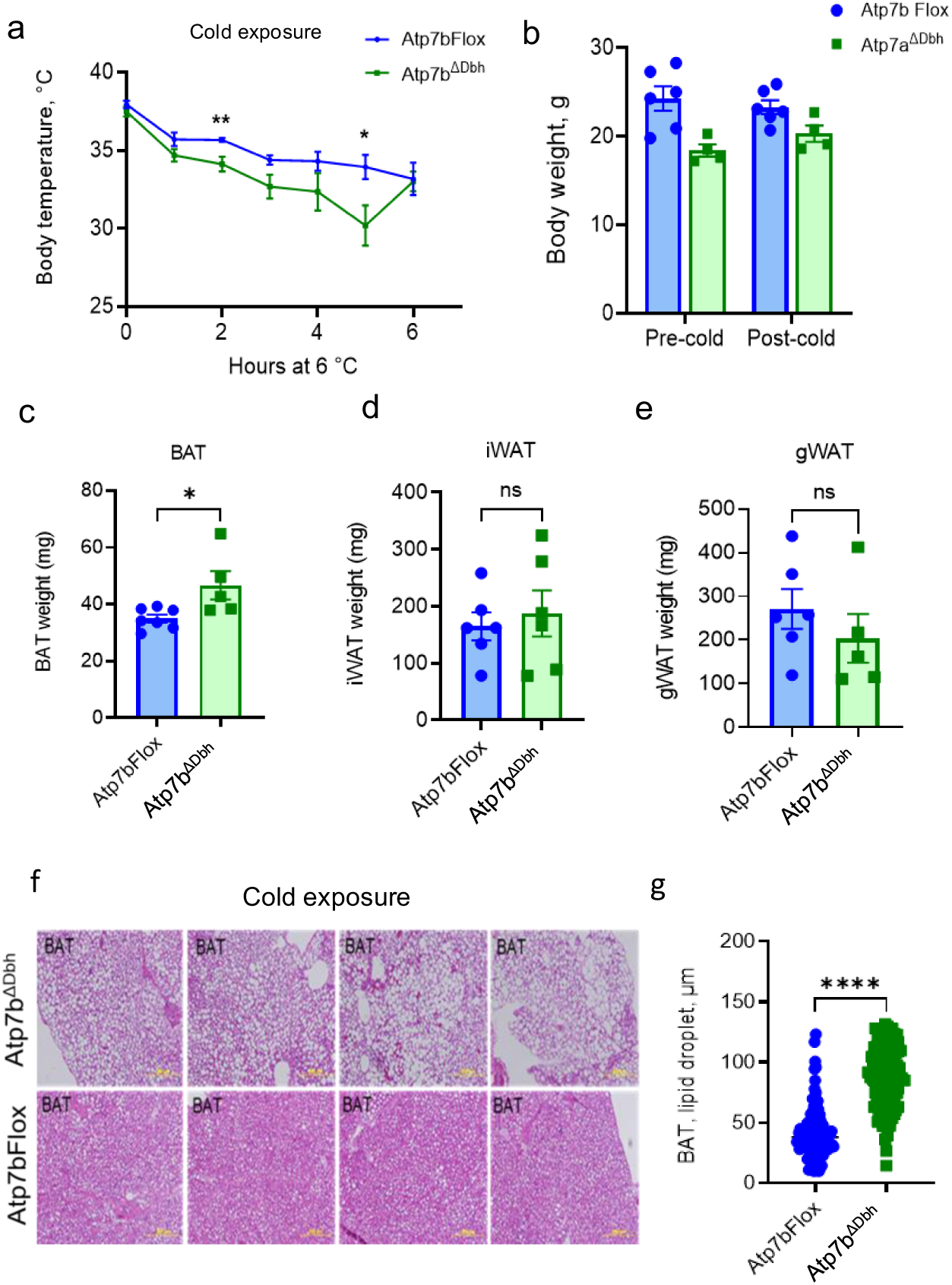
Impaired lipolysis in adipose tissues of *Atp7b^ΔDbh^* mice following acute cold stress. (**a**) Twelve-week-old male *Atp7b^ΔDbh^* and *Atp7bFlox* animals (n = 4 per genotype) were exposed to cold exposure (6°C) for 6 hours and showed a mild thermoregulatory defect. Although partial recovery was observed at 6 hours in both groups, *Atp7b^ΔDbh^* mice continued to trend toward hypothermia compared to *Atp7bFlox* controls. (**b**) The body weight of *Atp7b^ΔDbh^* and *Atp7bFlox* controls remained unchanged before and after cold exposure. (**c**) *Atp7b^ΔDbh^* animals displayed significantly increased BAT weight relative to controls. (**d, e**) gWAT and iWAT weights showed no significant difference between *Atp7b^ΔDbh^* and *Atp7bFlox* controls. (**f**) Representative H&E-stained BAT sections from *Atp7aFlox* and *Atp7a^ΔDbh^* mice following cold stress at 6°C for 6 hours revealed enlarged adipocytes in *Atp7a^ΔDbh^* animals, consistent with impaired lipolysis. (**g**) Quantification confirms significant increases in adipocyte size in BAT in *Atp7a^ΔDbh^* animals compared to the *Atp7bFlox* controls. Data are shown as mean ± SEM (*n* = 4 mice per group). Statistical significance was determined using Graphpad Prism Version 10, using an unpaired two-tailed Student’s *t*-test with Welch’s correction. \**p* < 0.05, **p < 0.01, ****p < 0.0001; ns = not significant.

## DISCUSSION

This study uncovered the specific roles of Atp7a and Atp7b in the structural and functional integrity of NE neurons. Atp7a is essential for the survival and function of these neurons, enabling NE biosynthesis, maintaining local metal homeostasis, and coordinating thermogenic and metabolic responses. In contrast, Atp7b plays a more subtle modulatory role, reflected in its milder impact on the NE:DA ratio and thermogenesis. However, the clear thermogenic phenotype in *Atp7b^ΔDbh^* mice, even in the presence of Atp7a, underscores a non-redundant, specialized role for Atp7b within catecholaminergic circuits. In both knockout strains, loss of one Cu transporter fails to induce compensatory upregulation of the other, and functional substitution does not occur. Notably, both *Atp7a^ΔDbh^* and *Atp7b^ΔDbh^* mice exhibit decreased Dbh protein abundance, indicating that DBH is vulnerable to Cu dyshomeostasis.

Our results provide a mechanistic insight into MD patients’ often-reported hypothermia (Fujisawa et al., 2022). We demonstrate that the loss of ATP7A function in NE neurons alone impairs thermoregulation. By linking Cu-dependent NE biosynthesis to systemic energy balance, our findings reframe MD hypothermia as a circuit-level pathology that occurs even when Cu status of peripheral tissues is unaffected (or restored). Additionally, our results highlight potential contributions of noradrenergic Cu misbalance to metabolic and neurologic phenotypes in neurodegenerative disorders involving Cu and/or catecholaminergic dysfunction. Our findings may help elucidate the role of LC neurons in driving metabolic phenotypes in Wilson disease, providing a framework to dissect central versus peripheral contributions to copper-related metabolic dysfunction.

Thermogenic neural circuitry comprises three components: thermosensory afferents, central processing regions, and sympathetic efferent outputs. Because both *Atp7a^ΔDbh^*and *Atp7b^ΔDbh^* mice exhibit warmth-seeking behavior associated with hypothermia, their thermosensory inputs appear intact. In contrast, both central and sympathetic efferent outputs are malfunctioning, especially in *Atp7a^ΔDbh^*mice. This is evidenced by hyperphagia, indicative of abnormal NE signaling to hypothalamic nuclei (Sayar-Atasoy et al., 2023; Sciolino et al., 2022), neuronal loss, and proteomic changes. High fat content in the BAT of both knockout strains before and after cold exposure is consistent with a deficit in NE signaling by sympathetic neurons. Taken together, our data identify a Cu-modulated noradrenergic pathway linking brainstem activity to peripheral metabolic responses. Within this framework, Cu emerges as an essential determinant of brain-body communication. This integrative paradigm illustrates that trace metal transport can directly shape neural network function and systemic adaptation, with broad implications for metabolic regulation and neurodegenerative disease pathogenesis.

## MATERIALS AND METHODS

### Animal studies

All animal experiments were conducted based on ARRIVE guidelines and followed protocols approved by the Johns Hopkins University Animal Care and Use Committee (ACUC; Protocol No. M023M337). Mice were housed in the Johns Hopkins Animal Care Facilities under specific pathogen-free (SPF) conditions with a controlled temperature of 22-24 °C, relative humidity of 40–60%, and a 14-hour light/10-hour dark cycle. Animals were group-housed (up to five per cage) in individually ventilated cages containing corn cob bedding or paper nesting material. Standard rodent chow (Teklad 2018 or Picolab5V75) and water were available ad libitum. Static cages were changed once per week, while ventilated cages were changed every two weeks. All animals were monitored daily by trained animal care staff for their general health, grooming, and activity levels. All animals used in this study were on a C57BL/6J (B6) genetic background. *Atp7a^LoxP/LoxP^* mice and *Atp7b^LoxP/LoxP^* mice were described previously in (Xiao et al., 2018) and (Muchenditsi et al., 2017), respectively.

To generate mice with a targeted deletion of *Atp7a* or *Atp7b* in noradrenergic cells, female *Atp7a^LoxP/LoxP^* or *Atp7b^LoxP/LoxP^* animals were crossed with males expressing Cre recombinase under the control of the dopamine β-hydroxylase (*Dbh*) promoter (Dbh-Cre; Jackson Laboratory strain B6.Cg-*Dbh^tm3.2(cre)Pjen^*/J). Genotypes were confirmed by PCR analysis of genomic DNA isolated from tail biopsies for the presence of *Cre*, *Atp7a^LoxP/LoxP,^* or *Atp7b^LoxP/LoxP^* alleles (**Fig. S1**). Primer sequences are listed in **Table S3**. All experiments were performed using age- and sex-matched 4- to 20-week-old adult male mice. Age-matched littermates lacking Cre served as the control group. Wild-type, *Atp7a^LoxP/LoxP^* or *Atp7b^LoxP/LoxP^*age-matched male mice showed no significant differences in Atp7a, Atp7b, or Dbh expression in the locus coeruleus, nor in tissue Cu levels. Therefore, wild-type and *LoxP/LoxP* (*Atp7a^LoxP/LoxP,^* or *Atp7b^LoxP/LoxP^)* animals were used interchangeably as controls in subsequent experiments unless otherwise specified. Only animals confirmed by PCR to have the correct genotype and within the designated age range were included in the study. No animals were excluded after the inclusion criteria were met, unless humane endpoints were reached. Allocation was based on genotype, and blinding was applied during data collection and analysis where possible.

### Copper measurements

Cu levels were measured from the freshly dissected mouse brain using Atomic Absorption Spectroscopy (AAS) (Perkin Elmer Pinnacle 900T). Approximately 30mg of tissue washed and perfused in 1X Phosphate Buffered Solution (PBS) was digested in 200 μl of trace element-grade nitric acid (HNO3) (Fisher Scientific, USA) for 2 hours at 95°C in metal-free 15 ml tubes (Labcon). The digested sample was further diluted using MilliQ water to adjust the nitric acid concentration to less than 0.2%. Metal content was quantified using calibration standards 20 ppb and 10, 20, 50, and 100 ppb and divided by tissue weight.

### Enzyme-linked immunosorbent assay (ELISA)

Serum norepinephrine levels were measured using NA/NE(Noradrenaline/Norepinephrine) ELISA Kit (Elabscience; E-EL-0047) following the manufacturer’s protocol. The color intensity was measured at 450 nm using a BMG Labtech Microplate reader. The concentration of norepinephrine in the samples was determined by comparing the OD of the samples to the standard curve. A reference standard curve was prepared using concentrations of 0, 0.31, 0.63, 1.25, 2.5, 5.0, 10.0, and 20.0 ng/mL

### Catecholamine analysis using mass spectrometry

Serum and brain concentrations of dopamine, norepinephrine, epinephrine, and serotonin were quantified by liquid chromatography-mass-spectrometry (LC–MS) following protein precipitation with methanol. For serum, 50 µL was extracted with 150 µL methanol; brain tissue was homogenized in 1 mL methanol with ceramic beads, and both were spiked with 5 µL of 1 µg mL⁻¹ dopamine-d₄ as internal standard. Supernatants were centrifuged (4 °C, 45 min), dried under vacuum, and reconstituted in 50 µL water. Calibration standards (0.2–200 ng mL⁻¹) were prepared from authentic stocks. LC separation was performed on an Agilent 1290 Infinity II UHPLC with a Waters Acquity BEH C18 column (2.1 × 100 mm, 1.7 µm) at 45 °C, using water with 0.1% formic acid (A) and methanol (B) at 0.40 mL min⁻¹, with a gradient of 2–98% B over 4 min.

### Comprehensive Laboratory Animal Monitoring System (CLAMS) analysis

Whole-body energy expenditure and metabolic parameters were assessed by indirect calorimetry using the Comprehensive Lab Animal Monitoring System (CLAMS; Columbus Instruments) at the Center for Metabolism and Obesity Research, Johns Hopkins University (Baltimore, MD, USA). The measurements included oxygen consumption (VO₂), carbon dioxide production (VCO₂), respiratory exchange ratio (RER), and energy expenditure (EE). Food and water intake, as well as physical activity, were also recorded.

Age-matched adult male mice (16 weeks old) (Wild-type; n=8, *Atp7a^ΔDbh^*; n=12, *Atp7b^ΔDbh^*; n=4) were individually housed in chambers under standard environmental conditions (22–24°C; 12-hour light/dark cycle) with ad libitum access to food and water. Mice were acclimated to the chambers for 24 hrs before data collection. Measurements were obtained over 48 hrs and analyzed across light and dark phases. Ambulatory activity was quantified based on consecutive X-axis infrared beam breaks. Energy expenditure was calculated using the Weir equation: EE (kcal/day) = VO2× [3.815 + (1.232×RER)]. To account for variation in body composition, energy expenditure, and metabolic parameters were normalized to both total and lean body mass (Sarver et al., 2020).

### Glucose Tolerance Test (GTT)

To assess systemic glucose handling, male mice (*Atp7a^ΔDbh^*, n=12; wild-type littermates, n=8; *Atp7b^ΔDbh^*, n=4) were fasted for 6 h before testing. D-glucose (1 mg/g body weight) was administered intraperitoneally, and blood glucose concentrations were determined at baseline (0 min) and at 15-, 30-, 45-, 60-, 90-, and 120-min post-injection using a handheld glucometer (NovaMax glucometer). For each time point, blood was obtained from awake mice by tail snipping on Novamax glucose test strips.

### Insulin Tolerance Test (ITT)

To evaluate insulin sensitivity, the same cohorts of male mice (*Atp7a^ΔDbh^*, n=12; wild-type, n=8; *Atp7b^ΔDbh^*, n=4) were fasted for 2 h before testing. Human recombinant insulin (1.0 U/g body weight; intraperitoneal) was administered, and blood glucose was measured at 0, 15, 30, 45, 60, 90, and 120 min using Novamax glucometer using Novamax glucose test strips. The blood was collected onto the strips by tail snipping on awake mice.

### Pyruvate Tolerance Test (PTT)

To assess hepatic gluconeogenic capacity, male mice (*Atp7a^ΔDbh^*, n = 12; wild-type, n = 8; *Atp7b^ΔDbh^*, n = 4) were fasted for 6 h, followed by intraperitoneal injection of sodium pyruvate (1–2 mg/g body weight). Blood glucose concentrations were monitored at baseline and 15, 30, 45, 60, 90, and 120-min post-injection using an Accu-Chek glucometer. Retro-orbital sampling was used for blood collection.

### Thermoneutral Housing

To assess metabolic responses under thermoneutral conditions, 10-week-old male mice, *Atp7aFlox*(n=5) and *Atp7a^ΔDbh^* (n=5) were housed at 30°C in a static cage with a filtered top in a temperature-controlled environmental chamber with a 12-hour light/dark cycle and ad libitum access to food and water. Mice were acclimated for 10 days before physiological or metabolic measurements to ensure full adaptation to thermoneutrality, the ambient temperature at which thermogenic demand is minimized.

### Cold Tolerance Test

Cold-induced thermogenesis was evaluated by exposing mice to acute cold stress following acclimatization in thermoneutral conditions. Age-matched (10 weeks) adult male mice *Atp7aFlox* (5), *Atp7bFlox* (n=6), *Atp7a^ΔDbh^* (n=5), and *Atp7b^ΔDbh^* (n=4) were individually housed at 6°C in cages without nesting material and with minimal bedding. Access to food was restricted during acute cold exposure. Core body temperature was measured using a rectal probe (Physitemp RET-3 Rectal Probe; Cat. No. NC9713069) at baseline (room temperature) and at 1-hour time intervals for up to 6 hours following cold exposure. Mice had free access to water throughout the experiment. The cold exposure was terminated earlier if a mouse’s core temperature dropped below 30°C, at which point the animal was euthanized by cervical dislocation to prevent hypothermic distress. Following cold exposure, animals were sacrificed, and tissues were harvested for downstream analyses.

### Haematoxylin and Eosin (H&E) staining and histology analysis

The brown adipose tissue (BAT), inguinal white adipose tissue (iWAT), and gonadal adipose tissue (gWAT) isolated from mice were fixed overnight in 10% formalin and embedded in paraffin wax as previously described(Lee et al., 2012). Sections (8 μm) were stained with hematoxylin and eosin at the Johns Hopkins University Reference Histology Core, Baltimore, MD, USA. Images were acquired using an Olympus light color microscope (Johns Hopkins University, Microscope Facility, Baltimore, MD, USA). ImageJ was used for the quantification of adipocyte diameters from H&E-stained sections of iWAT, gWAT, and BAT collected from 4 mice in each group (Wild type, *Atp7aFlox*, *Atp7bFlox*, *Atp7a^ΔDbh^*, and *Atp7b^ΔDbh^*).

### Membrane Protein Extraction and Immunoblotting

Whole mouse brains (∼450 mg) were homogenized in ice-cold homogenization buffer (3 ml) containing 0.25 M sucrose, 10 mM Tris-HCl (pH 7.5), and 1 mM EDTA, supplemented with a protease inhibitor cocktail (Roche, Basel, Switzerland) and 30 μl of each phosphatase inhibitor cocktails (Phosphatase inhibitor cocktail 2; Cat. No. P5726-1ML and Phosphatase inhibitor cocktail 3, Cat. No. P0044-1ML; Sigma-Aldrich). Homogenization was performed using a glass Dounce homogenizer with 50 strokes on ice. The homogenate was centrifuged at 100,000 × g for 20 minutes at 4°C using a Beckman Coulter benchtop ultracentrifuge. The pellet was resuspended in 100 μl homogenization buffer containing 0.1% Triton X-100. Protein concentrations were determined using the BCA assay (Pierce BCA Kit; Cat. No. 23227; Thermo Fisher Scientific) following the manufacturer’s instructions. Equal amounts of protein (15 µg) were resolved by 10% SDS-PAGE and transferred to PVDF membranes using semi-dry transfer in Tris-Buffered Saline transfer buffer for 10 min (Bio-Rad, California, USA). Immunoblotting was performed using the appropriate primary and secondary antibodies (**Table S4**). The PVDF membrane was blocked using blocking buffer (5% Milk in 1X TBST). The primary antibodies were diluted in 3% Milk in 1X TBST and incubated overnight at 4°C. The secondary antibodies were diluted in 3% Milk in 1X TBST and incubated for 1 h at room temperature. Relative expression levels of target proteins were quantified by densitometry using ImageJ software and normalized to the corresponding loading control.

### RNA Extraction and Quantitative PCR

Total RNA was extracted from fresh-frozen BAT, iWAT, and gWAT using TRIzol reagent (Life Technologies, Cat. No. 15596026), following the manufacturer’s protocol. RNA concentration and purity were assessed by 260/280 nm ratio using a BMG Labtech LVi plate reader. Reverse transcription was performed using the HiScript® IV RT supermix for qPCR with gDNA wiper (Vazyme, Catalog No. R423) to synthesize cDNA. Quantitative PCR (qPCR) was performed using the SupRealQ Ultra-HSYBR qPCR Master Mix (Vazyme) on a QuantStudio 6 Flex (Applied Biosystems) real-time PCR system. Target gene’s transcript levels were normalized to *Gapdh* mRNA, and relative transcript abundance was calculated using the comparative Ct (ΔΔCt) method(Livak and Schmittgen, 2001).

### Laser Ablation-Inductively Coupled Plasma-Time of Flight-Mass Spectrometry (LA-ICP-TOF-MS)

Spatial distribution of metals in 4% PFA fixed coronal brain sections was determined using LA-ICP-TOF-MS. Fresh-frozen brains from 20-week-old male mice were cryosectioned at 10 µm and mounted on Superfrost PLUS slides. Frozen sections (prepared as described below) were air-dried and ablated using an ImageBIO 266 laser ablation system (266 nm laser, 10 µm spot size, 200 Hz, 1.8–2.3 J/cm²) coupled to a TOFWERK ICPTOF S2 mass spectrometer. Helium was used as the carrier gas, and daily tuning was performed with NIST SRM612 glass. Elemental images for ^63^Cu, ^56^Fe, and ^66^Zn were acquired using TofPilot, calibrated against gelatin standards in Iolite (v4.8.6), and co-registered with confocal images to define regions of interest in the locus coeruleus.

### Immunofluorescence and confocal microscopy

Mice were anesthetized with 5% isoflurane and transcardially perfused with 1× sterile phosphate-buffered saline (PBS; 0.1 M, pH 7.4). Following perfusion, animals were sacrificed by cervical dislocation. Brains and adrenal glands were promptly dissected and post-fixed in 4% (w/v) paraformaldehyde (PFA) in 0.1 M phosphate buffer (PB) for 12 hours at 4 °C. Tissues were placed in 30% (w/v) sucrose prepared in 0.1 M PB for a minimum of 48 hours until they sank. Subsequently, tissues were embedded in Optimal Cutting Temperature (OCT) compound and stored at –80 °C until sectioning. Frozen tissues were sectioned at a thickness of 10 µm using a Leica cryostat (Leica CM3050 S) and mounted on Superfrost Plus slides (Fisherbrand; Catalog No. 1255015) for immunofluorescence. Sections were blocked and permeabilized in a buffer containing 2% bovine serum albumin (BSA), 0.2% saponin, and 15% fetal bovine serum (FBS; HiMedia) for 1 hour at room temperature. Primary antibodies were applied overnight at 4 °C in blocking buffer, using the following dilutions: Anti-Dopamine β-Hydroxylase (1:500) Anti-Tyrosine Hydroxylase (1:300) Anti-ATP7B (1:200) Anti-ATP7A (1:50) Anti-Iba1 (1:300) Anti-GFAP (1:200). After washing, sections were incubated with the appropriate secondary antibodies conjugated to Alexa Fluor 488, Alexa Fluor 568, or Alexa Fluor 647 (1:500; Goat anti-Rabbit or Donkey anti-Sheep) for 1 hour at room temperature in the dark. Nuclei were counterstained using ProLong ™ Gold Antifade Mountant with DAPI (Thermo Fisher Scientific), and coverslips were applied. Images were acquired using a Zeiss LSM 800 confocal microscope and processed using ImageJ (NIH) software for analysis and visualization.

### Laser capture microdissection

Mouse brain tissue sections were prepared from TH-immunostained specimens sectioned at 30 μm thickness using a microtome and mounted on polyethylene naphthalate membrane glass slides (Carl Zeiss Microscopy GmbH, Jena, Germany). Sections were placed into the dish holder of the microscope, and laser capture microdissection (LCM) was performed using the PALM MicroBeam Ultraviolet Laser Microdissection system on the AxioObserver Z.1 system.

TH-positive locus coeruleus regions were identified and micro-dissected from both left and right hemispheres of each tissue section at 10× magnification while keeping laser power to a minimum. A total of four locus coeruleus samples were collected per animal (bilateral dissection from two sections per animal). Isolated tissue samples were directly collected in the cap of collection tubes (AdhesiveCap 200 opaque). Samples in collection tubes were stored at −80°C until further processing.

### Sample Processing and Protein Digestion

To release tissue samples from the adhesive caps, 30 μL of lysis buffer containing 1% n-dodecyl-β-D-maltoside (DDM), 600 mM guanidine HCl, 150 mM NaCl, 10 mM HEPES (pH 7.4), and 15 mM TCEP was added to each collection tube. Samples were resuspended and transferred to new tubes. Sonication was performed using a Branson Sonifier 250 (Branson Ultrasonics), followed by centrifugation at 14,000 × g for 1 min. For decrosslinking, samples were incubated at 95°C for 30 min, then centrifuged again at 14,000 × g for 1 min.

Proteins were reduced and alkylated with 10 mM Tris(2-carboxyethyl) phosphine hydrochloride and 40 mM chloroacetamide at room temperature (22–25°C) for 1 h. The proteins were then digested with Lys-C (Lysyl endopeptidase MS grade; Fujifilm Wako Pure Chemical Industries Co, Ltd) at a final concentration of 10 ng/μL and incubated at 37°C for 3 h. Subsequently, trypsin digestion was conducted by diluting the guanidine HCl concentration from 600 mM to 200 mM with 50 mM TEABC buffer, adding trypsin (sequencing grade modified trypsin; Promega) at a final concentration of 10 ng/μL, and incubating at 37°C overnight (15–18 h).

### TMT Labeling and Fractionation

The resulting peptides were desalted using C18 StageTips (3M Empore; 3M) and labeled with 18-plex TMT reagents according to the manufacturer’s instructions (Thermo Fisher Scientific). The labeling reaction was performed at room temperature for 1 h, followed by quenching with 1/10 volume of 1 M Tris–HCl (pH 8.0). The labeled peptides were pooled and pre fractionated into 6 fractions using strong cation exchange (SCX) Stage-Tips (3M Empore) with the following elution conditions: fraction 1 (50 mM ammonium acetate, 20% ACN, 0.5% FA), fraction 2 (75 mM ammonium acetate, 20% ACN, 0.5% FA), fraction 3 (125 mM ammonium acetate, 20% ACN, 0.5% FA), fraction 4 (200 mM ammonium acetate, 20% ACN, 0.5% FA), fraction 5 (300 mM ammonium acetate, 20% ACN, 0.5% FA), and fraction 6 (80% ACN, 5% ammonium hydroxide). Subsequently, the fractionated samples were vacuum dried using a SpeedVac (Thermo Fisher Scientific) and stored at −80°C until use.

### Mass Spectrometry analysis of proteomes

Mass spectrometry analysis was performed as previously described with minor modifications (22). Peptide samples were analyzed on an Orbitrap Fusion Lumos Tribrid Mass Spectrometer (Thermo Fisher Scientific) coupled to an Ultimate 3000 RSLCnano nanoflow liquid chromatography system (Thermo Fisher Scientific). Each fraction was reconstituted in 16 μL of 0.1% formic acid (FA), and 15 μL was injected onto a trap column (Acclaim PepMap 100, C18, 5 μm, 100 μm × 2 cm, nanoViper; Thermo Fisher Scientific) at a flow rate of 8 μL/min. Peptides were separated on an analytical column (Easy-Spray PepMap RSLC C18, 2 μm, 75 μm × 50 cm; Thermo Fisher Scientific) using a linear gradient of solvent B (0.1% FA in 95% acetonitrile) at 0.3 μL/min. The column was interfaced with an EASY-Spray ion source operated at ∼2.0 kV. Chromatographic separation was carried out over 120 min. Protein identification and quantification were performed using Proteome Discoverer (version 3.2.0.450; Thermo Fisher Scientific).

### Statistical analysis

Statistical analysis was conducted in Perseus (version 1.6.0.7) (Tyanova et al., 2016). Data normalization was achieved by dividing reporter ion intensities by median protein values, followed by log2 transformation and median centering across samples. Differential protein expressions were assessed by Student’s two-sample t-test, with proteins showing q < 0.05 considered significantly different. For volcano plot generation, q-values were calculated using Significance Analysis of Microarrays (SAM) with permutation-based false discovery rate estimation and an S0 parameter of 0.1(Tusher et al., 2001). Principal component analysis (PCA) was performed using MetaboAnalyst (Pang et al., 2022). Functional enrichment of differentially expressed proteins was analyzed with Enrichr (Kuleshov et al., 2016) and Ingenuity Pathway Analysis (ThermoFisher).

## Acknowledgments

We thank Dr. Martina Ralle (OHSU) for metal analysis, Dr. Betty Eipper for providing high-quality antibody, Mr. Benjamin Devenney for assistance with the initial animal work, and Dr. Dwight Bergles for helpful advice. CLAMS studies were made possible by the Johns Hopkins IBBS Core Coins award to SL.

## Author contributions

Conceptualization: SR and SL

Methodology: KM, AW, AC, AT, MP, CT, KG

Investigation: SR, YW, NS, KK, AM, SA, LTH

Visualization: SR, YW, NS, AC, SA, KK

Supervision: SL, CHN, AK, TO’H

Writing—original draft: SR, YW, NS, KM, SL

Writing—review & editing: SR, SA, AK, MP, TO’H, KG, SL

## Data and materials availability

The mass spectrometry proteomics data have been deposited to the ProteomeXchange Consortium via the PRIDE partner repository with the dataset identifier PXD069564 and 10.6019/PXD069564

## Supplemental Materials and Methods LA-ICP-TOF-MS

### a) Preparation and quantification of gelatin standards

Matrix-matched LA-ICP-ToF-MS gelatin standards were made by spiking a molten gelatin solution with multi-elemental stock solutions to convert counts from elements (m/z) from the LA-ICP-TOF-MS into concentrations (ppm). Briefly, 10% (w/v) porcine gelatin (Bloom 300, Sigma Aldrich, St. Louis, MO, USA) solution in ultrapure H2O kept above the melting temperature at 55 oC. Following gelatin dissolution (care was taken to mix thoroughly while introducing few bubbles via sonication), the first set of gelatin standards was spiked with IV-Stock-74434 (Inorganic Ventures, 1000μg/mL: As, B, Ca, Cd, Co, Cr, Cu, Fe, K, Mg, Mn, Mo, Na, Pb, S, Se, V, Zn) at concentrations of 0, 20, 35 and 50 ppm to calibrate concentrations of 63Cu, 56Fe and 66Zn. The second set of standards was prepared to evaluate Na/P at concentrations of 0, 2000, 4000, and 8000 ppm using monosodium phosphate (Reagent Plus grade, ≥99.0%, Millipore Sigma, St. Louis, MO, USA). Both sets of gelatin standards were maintained at 55 oC before pipetting 250μl from each gelatin standard directly on a pre-cooled cryotome chuck (−20 oC). Gelatin was allowed to freeze completely in the cryostat for at least 5 minutes before sectioning. Sectioning was accomplished using a Leica CM3050S cryostat (Leica Biosystems, DerPark, IL, USA) with a chamber temperature of −22 °C and objective temperature of −20 °C. Once solidified, standards and murine brain tissues were sectioned at 10 μm thickness and adhered to positively charged glass slides (Superfrost Plus, Thermo Fisher Scientific, Waltham, MA, USA). The elemental concentrations of the gelatin standards were confirmed with ICP-QQQ-MS and ICP-OES as previously described (21).

### b) Data acquisition and analysis

Tissue samples from 20-week-old male mice (n=4) and standards mounted on glass slides were ablated using the ImageBIO 266 laser ablation system (Elemental Scientific Lasers, Bozeman, MT, USA), which is equipped with a 266 nm laser, an ultra-fast, low-dispersion TwoVol3 ablation chamber, and a dual concentric injector. The aerosolized sample was transferred to the TOFWERK icpTOF S2 mass spectrometer (TOFWERK AG, Thune, Switzerland), where the elemental content was analysed in real time according to mass/charge (m/z) ratio. Daily tuning was performed using the NIST SRM612 glass certified reference material (National Institute for Standards and Technology). High intensities for 140Ce and 55Mn were used to optimize torch alignment, lens voltages, and nebulizer gas flow while maintaining low oxide formation based on the 232Th16O+/232Th+ ratio (less than 0.5). Laser ablation sampling was performed at 50% laser power (no dosage) at a repetition rate of 200 Hz using a circular 10 μm spot size. Helium was used as the carrier gas (chamber and cup helium at 350 ml/min), and laser fluence was 1.80-2.26 J/cm2. Ten lines of gelatin standards were ablated using the same parameters but with the raster spacing set at 30 μm to ensure clean ablation of each raster. Scanning data was recorded using TofPilot (version 1.3.4.0, TOFWERK AG) and saved in the open-source Hierarchical Data Format 5 file format. Image generation and concentration calibration were conducted using the Iolite software package (version 4.8.6, Elemental Scientific Lasers). Finally, mass-calibrated laser images were overlayed with confocal images (using house-built software, unpublished) to locate the LC, where ROIs were made around LC to quantify 63Cu, 56Fe and 66Zn. 31P was used to normalize each element’s concentration.

### CATECHOLAMINE MEASUREMENTS

#### a) Sample Preparation for Serum

Serum concentrations of dopamine, norepinephrine, epinephrine, and serotonin were quantified by liquid chromatography–mass spectrometry (LC–MS) following protein precipitation. Briefly, 50 µL of serum was combined with 150 µL of methanol, vortexed to ensure complete protein precipitation, and centrifuged at 4 °C for 45 min at high speed. LC vials were prepared by spiking 5 µL of 1 µg mL⁻¹ dopamine-d₄ as an internal standard. A 50 µL aliquot of the supernatant was transferred to the spiked vials, and 10 µL was diluted with 490 µL of methanol. After vortexing, 50 µL of the diluted supernatant was transferred to a new vial. Additional dilution was performed when necessary to bring serotonin concentrations within the calibration range. All samples were dried under vacuum and reconstituted in 50 µL of water prior to analysis.

#### b) Sample Preparation for Brain Tissue

Brain tissue was homogenized using five 1.4 mm ceramic beads and 1 mL of methanol in a Bead Mill 4 Mini Homogenizer (speed 5, 30 s). Homogenates were centrifuged at 4 °C for 45 min. Supernatants were processed following the same procedure as serum samples: spiking with 5 µL of 1 µg mL⁻¹ dopamine-d₄, dilution with methanol, and additional dilution as needed to bring serotonin and dopamine within the calibration range. Samples were dried under vacuum and reconstituted in 50 µL of water prior to LC–MS analysis.

#### c) Calibration Standards

Calibration standards were prepared by serial dilution from 1 mg mL⁻¹ stock solutions of dopamine, norepinephrine, epinephrine, and serotonin. The calibration range for all analytes was 0.2–200 ng mL⁻¹.

#### d) Chromatographic Separation

Liquid chromatography was performed on an Agilent 1290 Infinity II UHPLC system (Agilent Technologies, Santa Clara, CA, USA) equipped with a Waters Acquity UPLC BEH C18 column (1.7 µm, 2.1 × 100 mm). The mobile phase consisted of water with 0.1% formic acid (A) and methanol (B). The gradient program was as follows: 2% B (0–1.0 min), linear increase to 98% B (1–4 min), hold at 98% B (4–6 min), return to 2% B (6–6.1 min), and re-equilibrate at 2% B (6.1–10 min). The flow rate was 0.40 mL min⁻¹, the column temperature was maintained at 45 °C, and the injection volume was 10 µL.

### LC-MS PROTEOMICS ANALYSIS OF LASER-MICRO-DISSECTED LC

#### a) Mass Spectrometry Detection

Mass spectrometric analysis was performed using a Sciex 6500+ triple quadrupole mass spectrometer (Sciex, Framingham, MA, USA) operating in positive electrospray ionization (ESI) mode. Multiple reaction monitoring (MRM) was used for detection and quantification of target compounds (Table 1). The declustering potential was 60 V, entrance potential 10 V, and collision cell exit potential 8 V. Source conditions were as follows: curtain gas, 35 psi; ion spray voltage, 5500 eV; source temperature, 500 °C; ion source gas 1, 65 psi; ion source gas 2, 55 psi. Quantification was performed using Sciex MultiQuant 3.1 software, based on the peak area ratio of the analyte to the internal standard.

#### b) Data acquisition

Calibration curves were constructed by plotting the peak area ratio of each analyte to the internal standard against the nominal concentration of calibration standards. Linear regression with 1/x weighting was applied to all calibration curves. Analyte concentrations in samples were calculated from the corresponding calibration curve. Method performance was evaluated by assessing linearity (R² ≥ 0.99). Limits of detection (LOD) and quantification (LOQ) were determined based on a signal-to-noise ratio of 3:1 and 10:1, respectively.

Mass spectrometric data acquisition was performed in data-dependent mode with full scan acquisition covering a mass-to-charge ratio (m/z) range of 300–1800 using “Top Speed” mode with a 3 s cycle time. Precursor ion scans (MS1) were acquired at the resolution of 120,000 at m/z 200, while fragment ion scans (MS2) were obtained using higher-energy collisional dissociation (HCD) at 35% collision energy and detected at the resolution of 50,000 at m/z 200. Automatic gain control was set to 1 × 10⁶ ions for MS1 and 5 × 10⁴ ions for MS2, with maximum injection times of 50 ms and 100 ms, respectively. Precursor isolation was performed using a 1.6 m/z window with 0.4 m/z offset. Dynamic exclusion was applied for 30 s, and singly charged precursors were excluded. Internal mass calibration was achieved using the lock mass function (m/z 445.12002) from ambient air.

#### c) Data Processing

The mass spectrometry data analysis was conducted as described previously with minor modifications. Protein identification and quantification were performed using Proteome Discoverer software (version 3.2.0.450; Thermo Fisher Scientific). For MS2 preprocessing, the top ten peaks within each 100 Da window were selected for database searching. Tandem mass spectra were searched against the UniProt mouse protein database (UP000000589, which included both Swiss-Prot and TrEMBL and was released in January 2019 with 55,435 entries), which contained protein entries with common contaminants (115 entries) using SEQUEST HT search algorithms.

Search parameters included: trypsin digestion with up to two missed cleavages allowed, precursor mass tolerance of 10 ppm, fragment mass tolerance of 0.02 Da, carbamidomethylation of cysteine (+57.02146 Da) and TMT labeling (+304.20715 Da) on lysine residues and peptide N-termini as fixed modifications, and methionine oxidation (+15.99492 Da) as a variable modification. The minimum peptide length was set to six amino acids, with at least one peptide required per protein identification.

Peptide-spectrum matches (PSMs) and proteins were filtered to achieve a 1% false discovery rate using Percolator and Protein FDR Validator nodes, respectively. For TMT quantification, reporter ion integration utilized the most confident centroid method with 20 ppm mass tolerance for MS2-level HCD fragmentation. Both unique and razor peptides contributed to protein quantification, with a co-isolation threshold of 50%. Signal-to-noise ratios determined reporter ion abundances, with missing values replaced by minimum intensity values and an average signal-to-noise threshold of 50. Protein grouping followed strict parsimony principles, and final protein abundances were calculated by summing all corresponding PSM reporter ion intensities.

#### d) Statistical Analysis

Statistical analysis of mass spectrometry data was conducted using Perseus software (version 1.6.0.7). Data normalization was performed by dividing the reporter ion intensities by the median protein values, followed by a log2 transformation and median centering of the relative abundance values for each sample. Differential protein expressions between comparison groups were assessed using Student’s two-sample t-test. Proteins exhibiting q-values < 0.05 were designated as significantly differentially expressed. For volcano plot generation, q-values were determined using Significance Analysis of Microarrays (SAM) with permutation-based false discovery rate estimation and an S0 parameter of 0.1. Principal Component Analysis (PCA) analysis was conducted using the MetaboAnalyst tool. Functional enrichment analysis was performed using Enrichr to identify significantly enriched biological pathways and Gene Ontology (GO) terms associated with differentially expressed proteins.

**Fig. S1.**
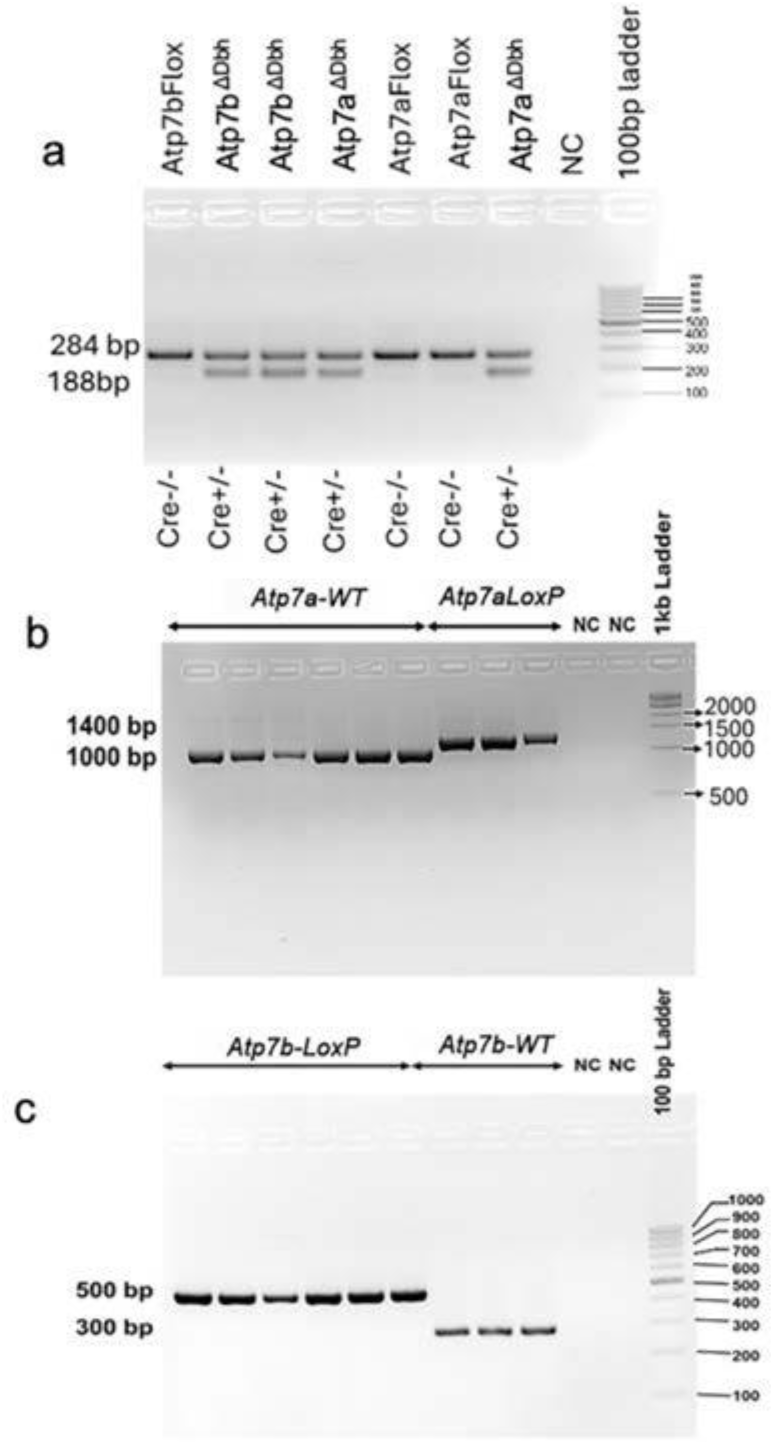
PCR validation of mouse genotypes. (a) Agarose gel electrophoresis (2% agarose in 1× Tris-acetate-EDTA buffer) showing PCR amplification of the Cre transgene. (b) PCR validation of Atp7aFlox alleles. (c) PCR validation of Atp7bFlox alleles.

**Fig S2:**
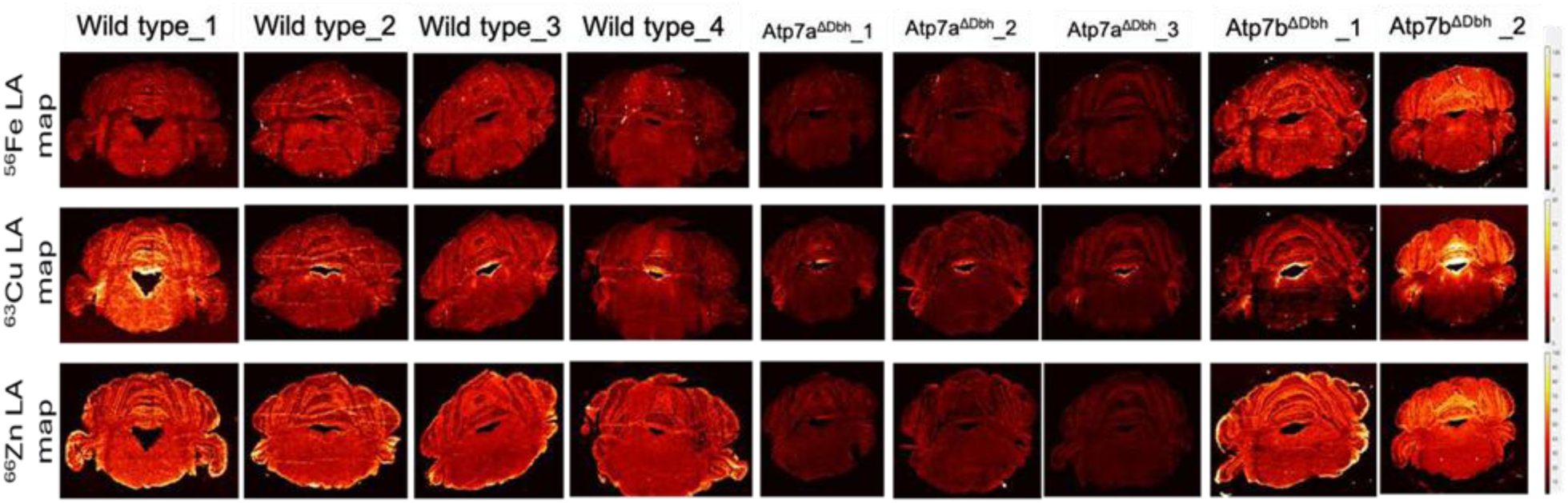
Metal maps for Cu, Zn, Fe and P on coronal brain sections from Wild type, Atp7a^ΔDbh^ and Atp7b^ΔDbh^ using LA-ICPMS

**Fig. S3.**
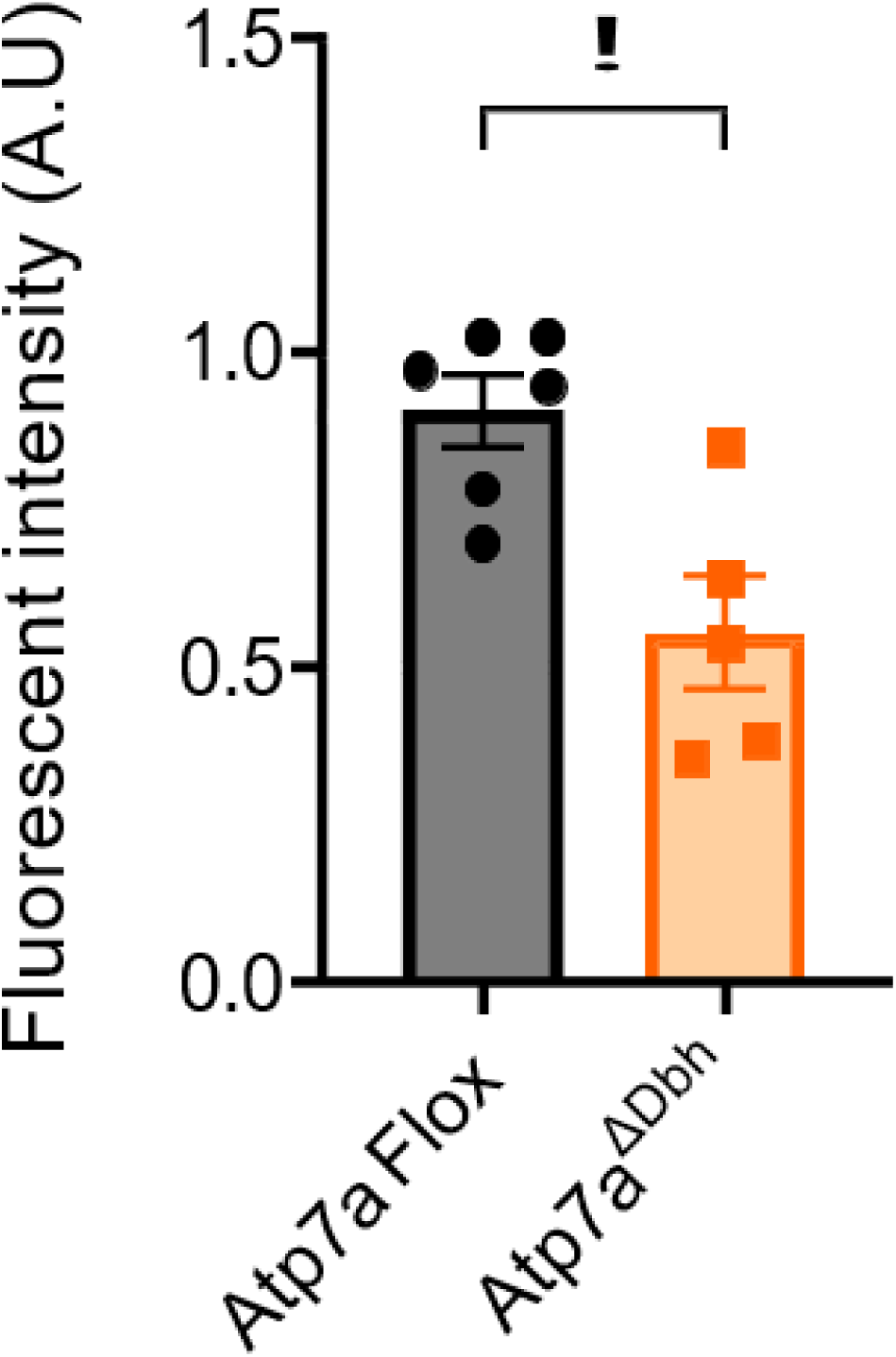
**(a)** Quantification of Atp7b fluorescent intensity in noradrenergic neurons of the Locus Coeruleus (LC), identified by TH staining, in *Atp7a^ΔDbh^* and *Atp7aFlox* coronal brain sections. n=3 animals were analyzed. Data were analyzed using GraphPad Prism 10 and are presented as mean ± SEM. Statistical significance was determined using an unpaired two-tailed Student’s t-test: *p* < 0.05, \**p* < 0.01, \*\**p* < 0.001.

**Fig. S4.**
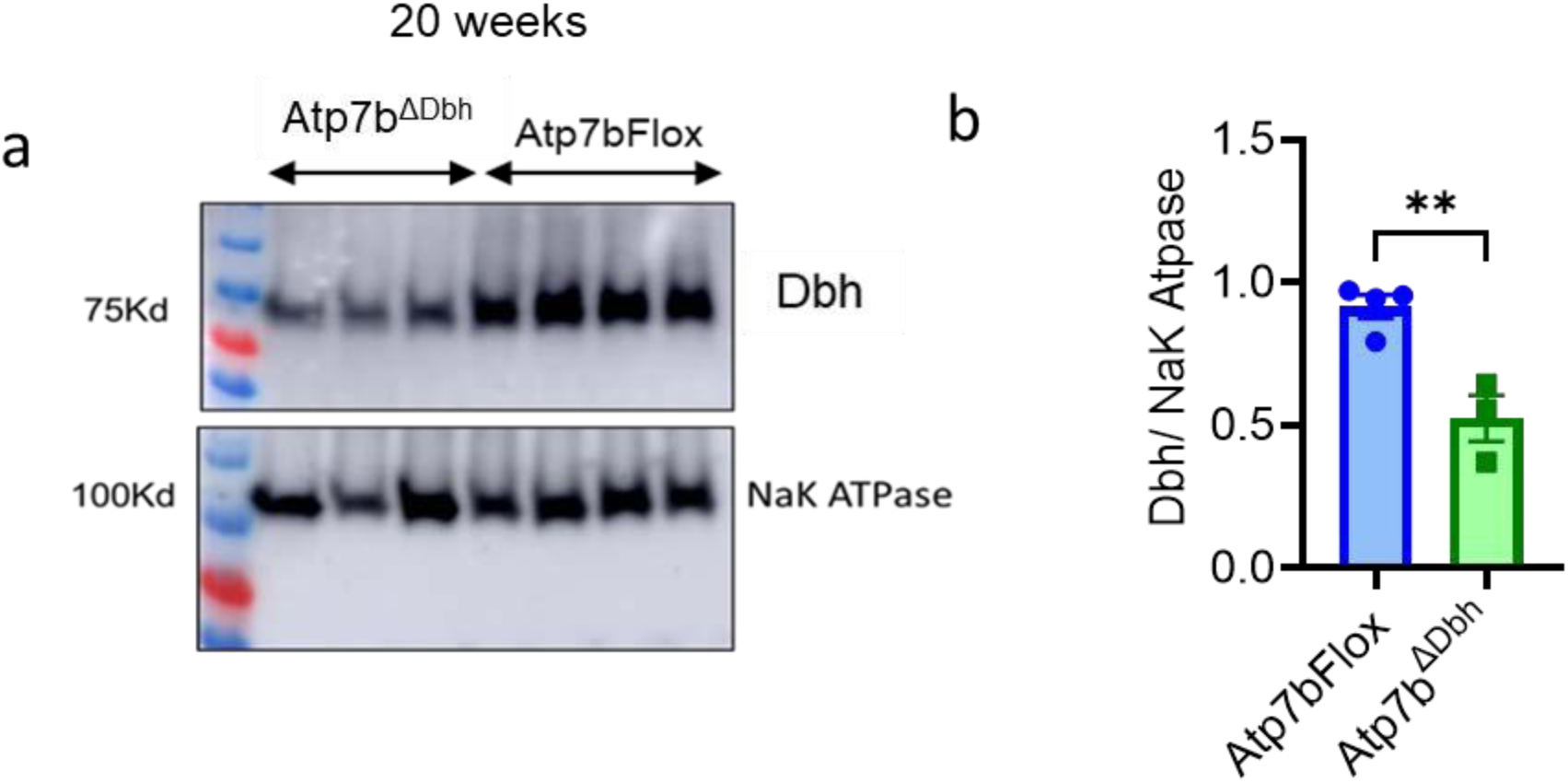
Immunoblot showing (a) Dbh from the membrane fraction isolated from whole brain homogenate from Atp7aFlox (n = 4) and Atp7b^ΔDbh^ (n=4) and Atp7bFlox (20 weeks) (b) Quantification of DBH normalized to Na+/K+ ATPase shows reduced expression in Atp7b^ΔDbh^ compared to the Atp7bFlox controls. Statistical significance was determined using GraphPad Prism version 10 software using an unpaired Student’s t-test with Welch’s correction. \**p* < 0.05, \*\**p* < 0.01, ns = not significant.

**Fig S5:**
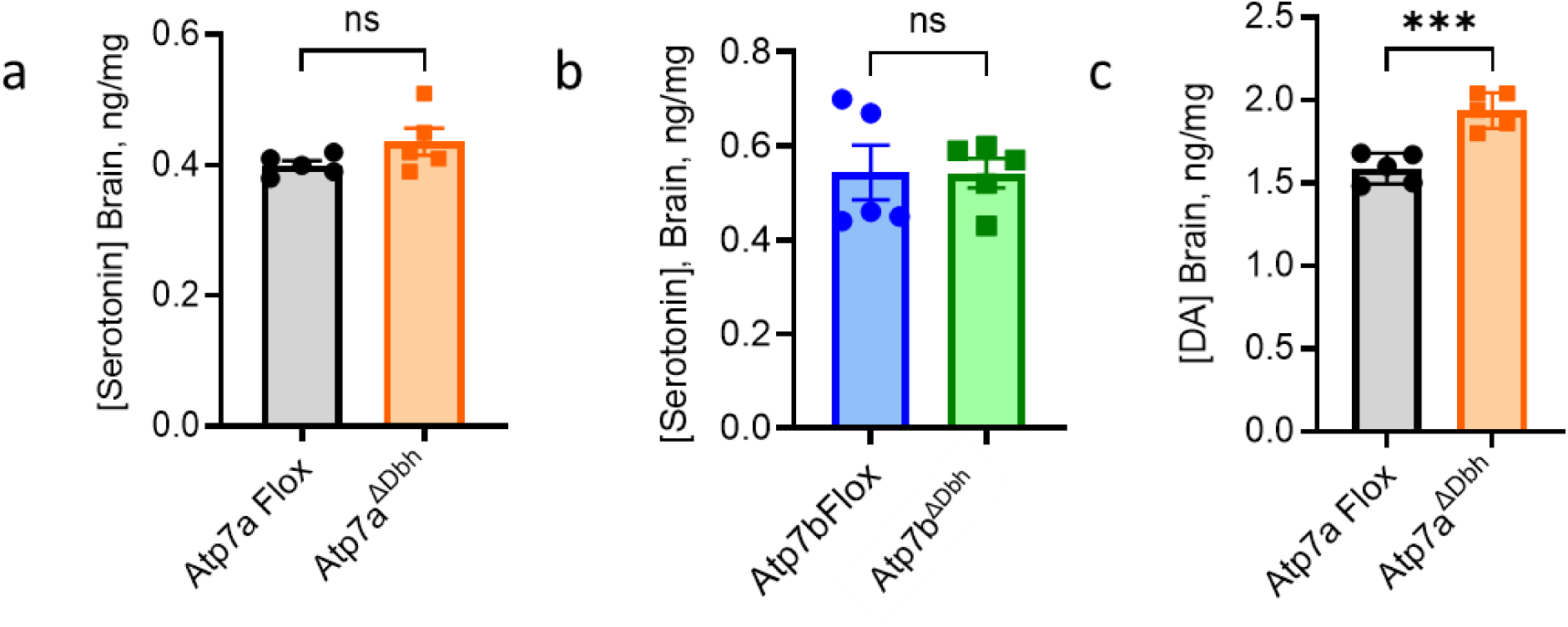
Inactivating copper transporters in Dbh neurons affects brain and serum Norepinephrine levels. (a, b) LC–MS analysis of whole-brain homogenates shows no significant change in serotonin levels in *Atp7a^ΔDbh^* (a) or *Atp7b^ΔDb^*^h^ (b) relative to *LoxP/LoxP* controls (20 weeks, n = 5). (c) Brain dopamine (DA) levels are significantly elevated in *Atp7a^ΔDbh^* mice compared to *Atp7aFlox*. Data are mean ± S.E.M. Statistical significance was determined using unpaired two-tailed Student’s t-test with Welch’s correction; *p < 0.05, **p < 0.01, *** p < 0.001.

**Fig S6:**
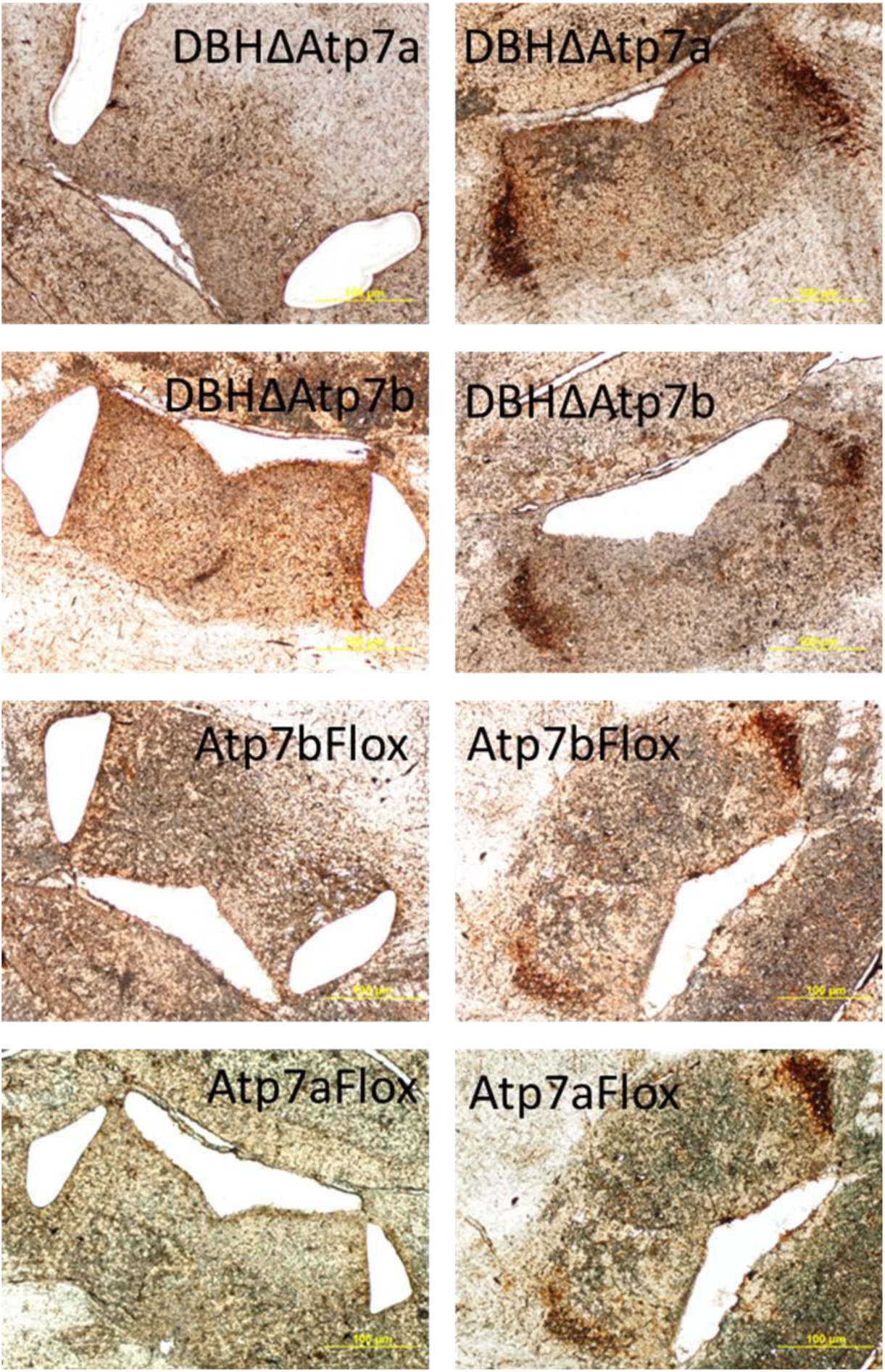
Immunohistochemistry was performed on 30 µm-thick coronal brain sections using a sheep anti-tyrosine hydroxylase antibody. Tyrosine hydroxylase-positive regions were identified and isolated using Zeiss Laser Capture Microdissection (LCM). The dissected tissues were subsequently subjected to LC-MS-based proteomic analysis.

**Fig S7:**
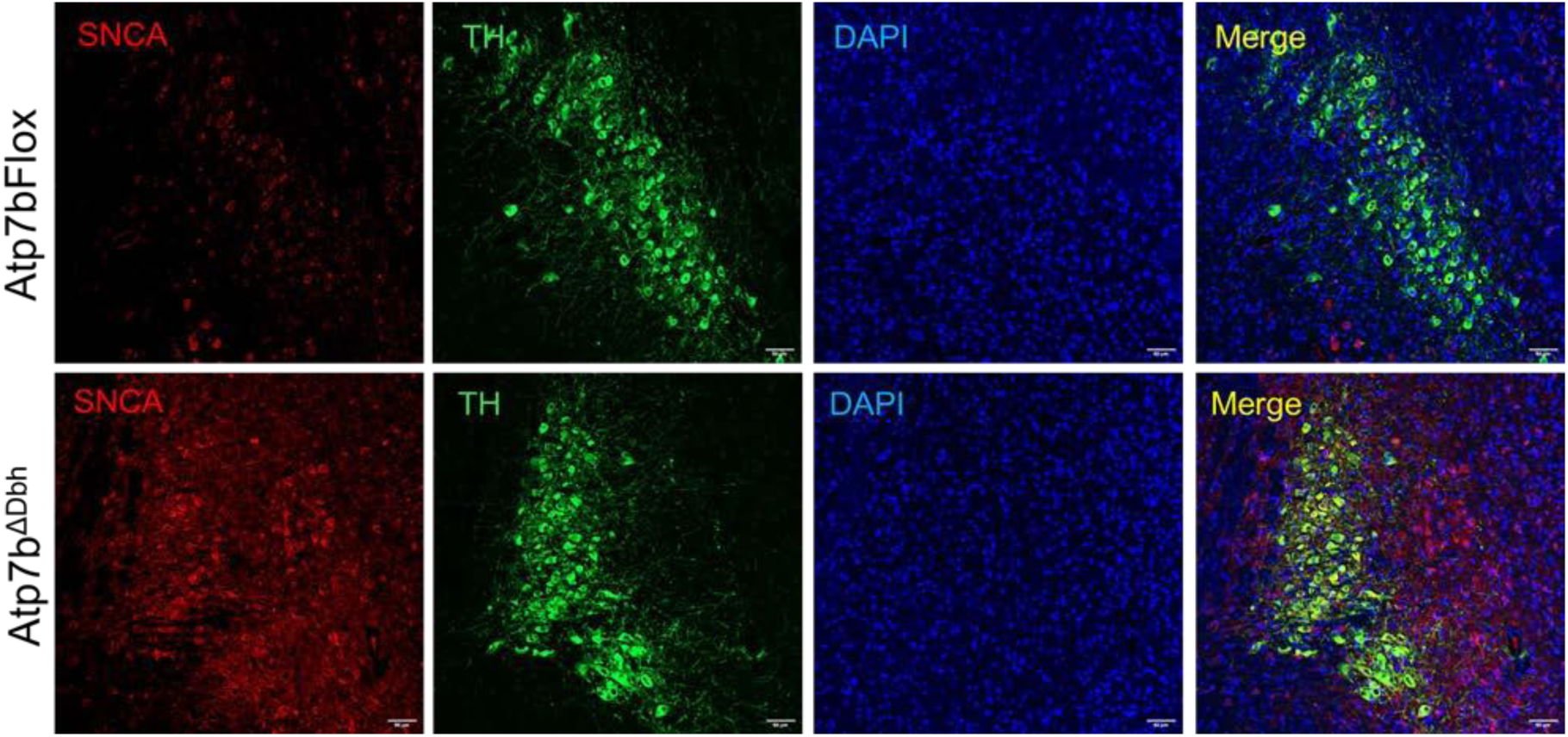
Immunostaining for SNCA (red) in Atp7b^ΔDbh^ shows upregulation of SNCA in TH+ve (green) noradrenergic neurons in the LC region compared to the Atp7bFlox controls (20 weeks)

**Fig S8:**
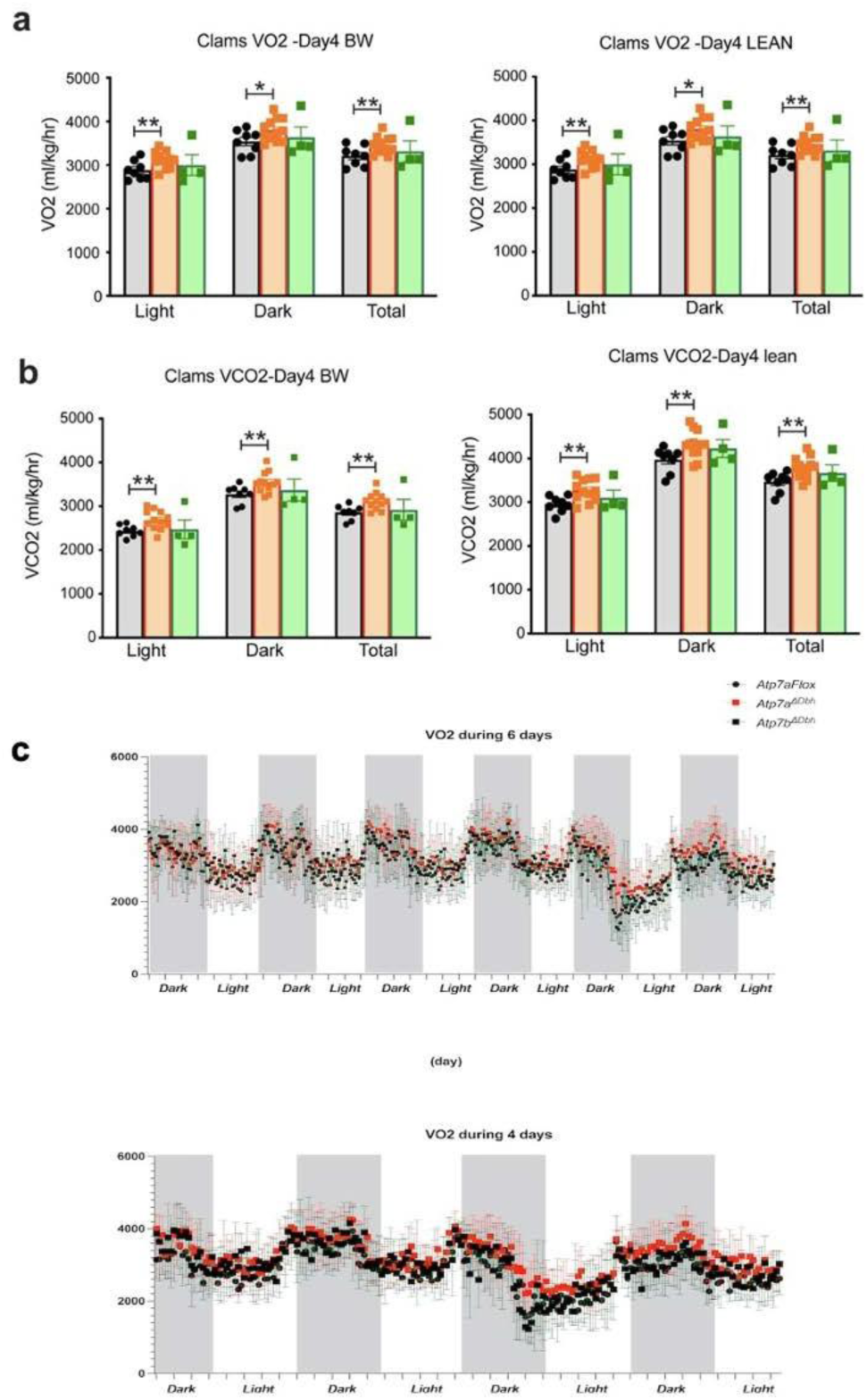
Conditional inactivation of Atp7a in Dbh neurons disrupts systemic energy metabolism. Whole-body metabolic parameters were measured in wild-type (black), Atp7a^ΔDbh^, and Atp7b^ΔDbh^ male mice using the Comprehensive Lab Animal Monitoring System (CLAMS; Columbus Instruments) over a continuous 48-h recording period following 24 h acclimatization. (a) Oxygen consumption (VO₂) and (b) carbon dioxide production (VCO₂) normalized to either lean mass or body weight. (c) Respiratory exchange ratio (RER). Light and dark phases are indicated, and whole-day averages are shown. Data represent mean ± SEM, with individual animals shown as dots. Statistical analyses were performed using two-way ANOVA followed by Tukey’s multiple comparison test; *p < 0.05, **p < 0.01, ***p < 0.001, ns = not significant

**Fig S9:**
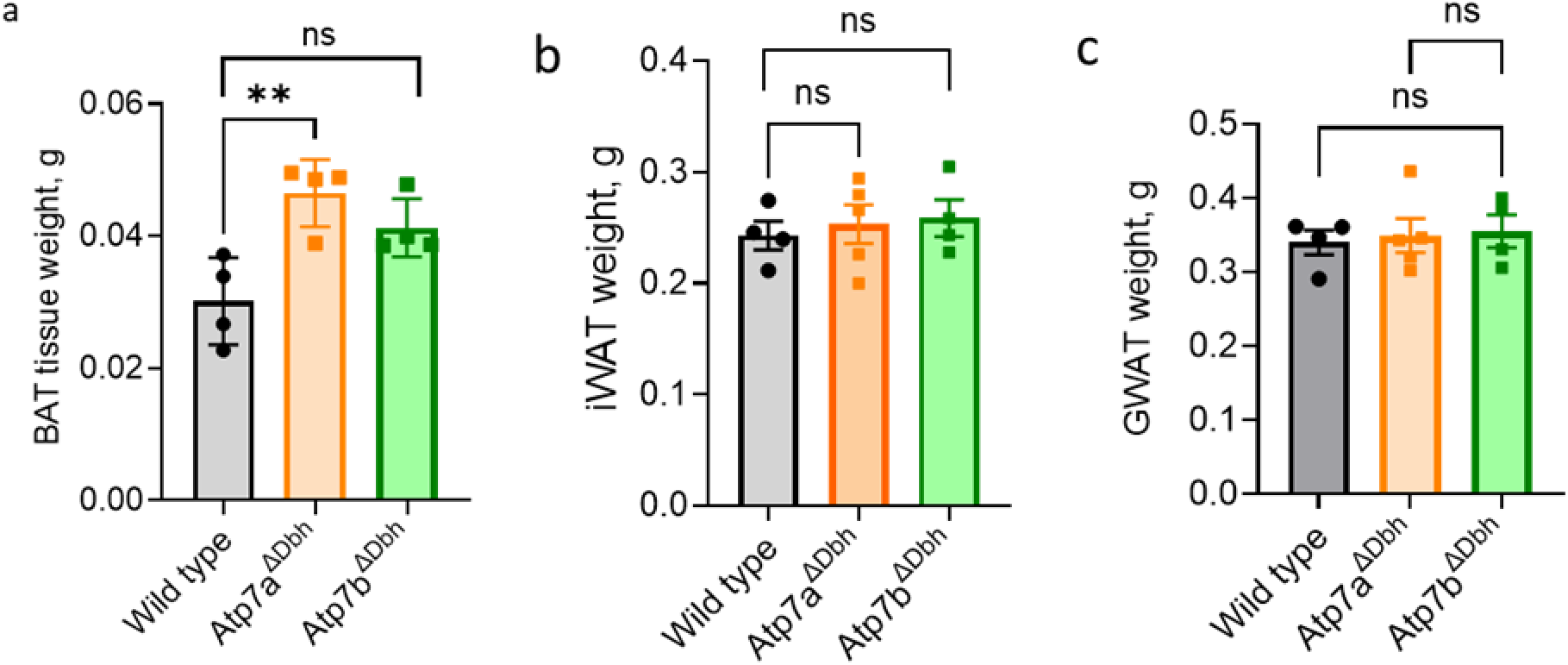
Weight of adipose tissues isolated from mice kept at room temperature (22-24⁰ C) (a) BAT from Atp7a^ΔDbh^ animals shows significantly increased weight compared to the Wild type (20 weeks). Atp7b^ΔDbh^ did not show any significant change (b) iWAT (c) gWAT did not reveal any significant difference in adipose tissue weight in mutants compared to the wild type controls. Data were represented as Mean ± SEM. Statistical significance was determined using GraphPad Prism version 10 software using an unpaired Student’s t-test with Welch’s correction. *p < 0.05, **p < 0.01, ***p < 0.001, ns = not significant.

**Fig S10:**
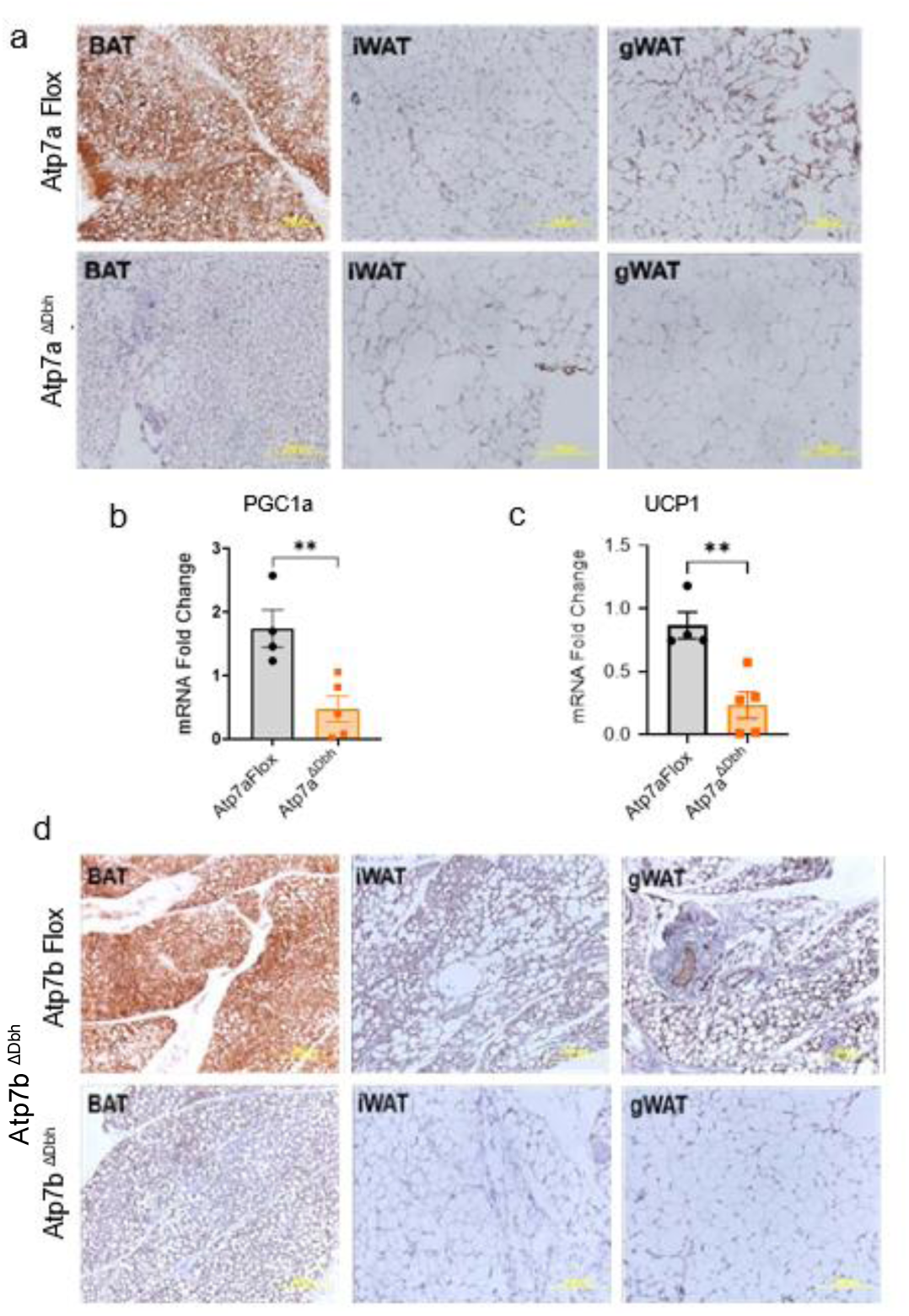
Downregulated thermogenic gene expression in adipose tissues in Atp7a^ΔDbh^ and Atp7b^ΔDbh^ mice following cold exposure. (a) Immunohistochemistry for UCP1 of adipose tissues, such as BAT, gWAT and iWAT shows intense DAB staining in Atp7aFlox animals while Atp7a^ΔDbh^ shows minimal staining suggesting reduced expression of UCP1 following acute cold stress; n=3. (b, c) Atp7a^ΔDbh^ mice (n=5) show reduced PGC1α and UCP1 mRNA expression in BAT compared to Atp7aFlox mice (n=4) normalized to GAPDH mRNA expression. (e) Immunohistochemistry for UCP1 of adipose tissues, such as BAT, gWAT and iWAT shows intense DAB staining in Atp7bFlox animals while Atp7b^ΔDbh^ shows minimal staining suggesting reduced expression of UCP1 following cold exposure; n=3). Statistical significance was determined using Graphpad Prism Version 9, using an unpaired two-tailed Student’s *t*-test with Welch correction; *P* < 0.05, *P* < 0.01.

**Fig S11:**
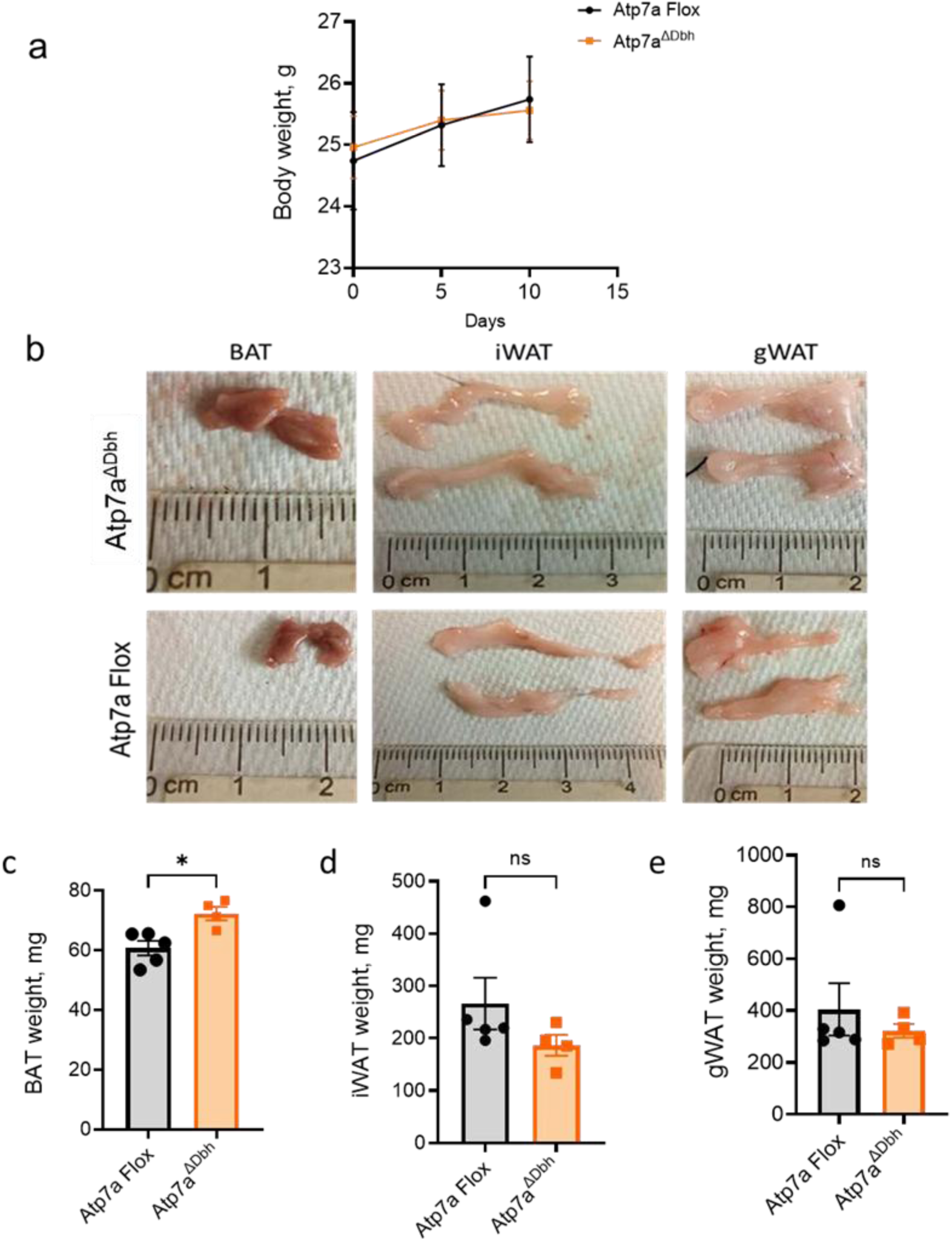
(a) Body weight of *Atp7a^ΔDbh^* kept at prolonged thermoneutrality (30 °C) shows a similar trend to that of *Atp7aFlox* (b) Adipose tissue morphology isolated from *Atp7a^ΔDbh^* and *Atp7aFlox* at thermoneutrality. Brown Adipose tissue (BAT) seems to be pale and enlarged. iWAT and gWAT seem to be comparable between *Atp7a^ΔDbh^* and *Atp7aFlox*. (c-e) Adipose tissue weight isolated from *Atp7a^ΔDbh^* and *Atp7aFlox* mice kept in prolonged thermoneutrality (30⁰C). Data were represented as Mean ± SEM. Statistical significance was determined using GraphPad Prism version 10 software using an unpaired Student’s t-test with Welch’s correction. \**p* < 0.05, \*\**p* < 0.01, \*\*\**p* < 0.001, ns = not significant.

**Fig S12:**
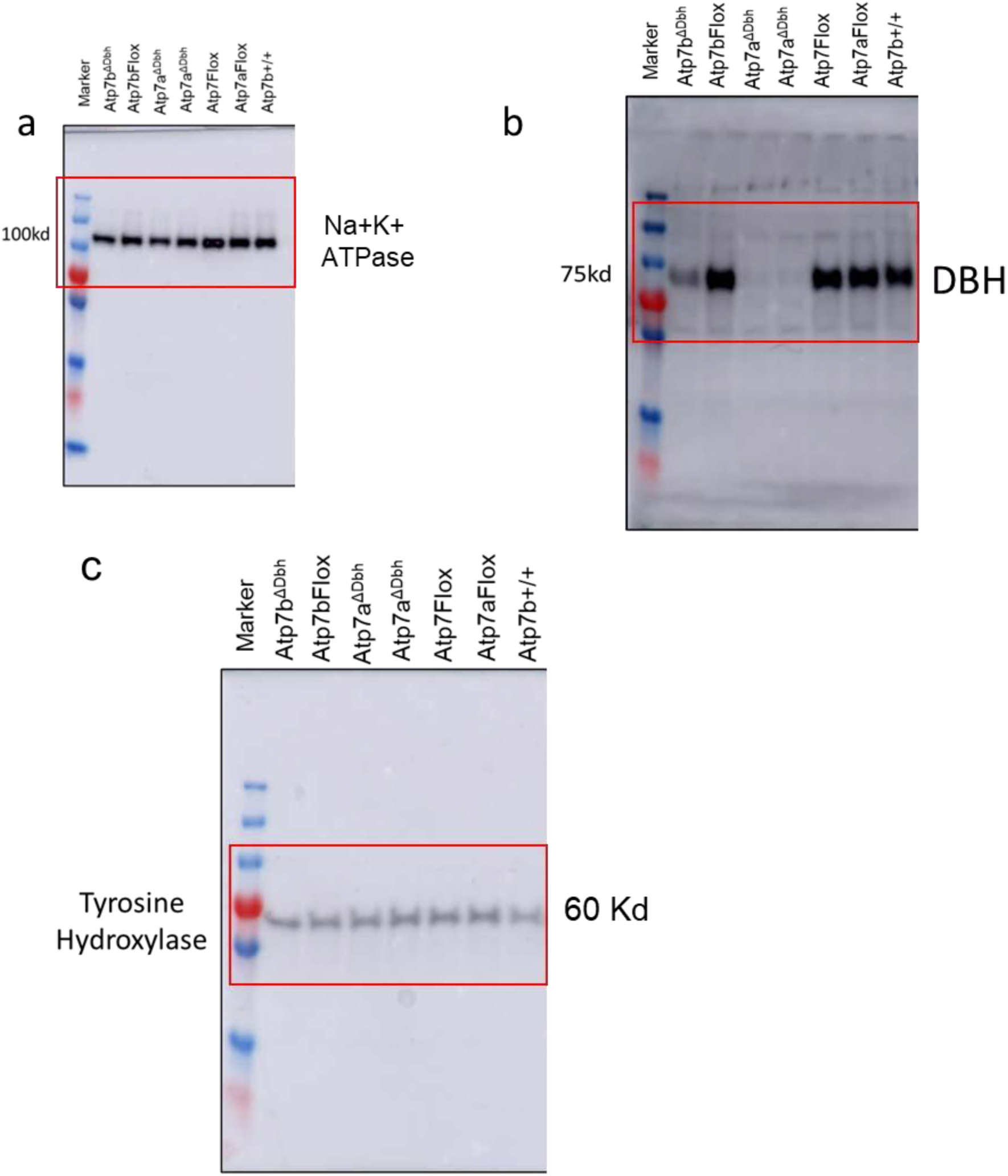
Uncropped immunoblot showing (a) Na^+^K^+^ATPase and (b) DBH from the membrane fraction from whole brain homogenate. (c) Immunoblot for TH from soluble and membrane fractions from whole brain homogenate.

**Table S1:** Elemental quantification at LC region in brain sections (copper is highlighted in blue)

| Genotype | Element | LC-Left | LC-Rright | LC AVG |
| --- | --- | --- | --- | --- |
| Wild type 1 | Fe56_ppm | 26.06644311 | 30.94934954 | 28.5079 |
|  | Cu63_ppm | 13.23363265 | 12.76365315 | 12.99864 |
|  | Zn66_ppm | 29.26250966 | 35.37463234 | 32.31857 |
| Wild type 2 | Fe56_ppm | 24.13445349 | 22.82077292 | 23.47761 |
|  | Cu63_ppm | 8.453597428 | 7.352046195 | 7.902822 |
|  | Zn66_ppm | 26.54174485 | 25.39997018 | 25.97086 |
| Wild type 3 | Fe56_ppm | 30.4119277 | 20.06864234 | 25.24029 |
|  | Cu63_ppm | 25.06456501 | 17.93140846 | 21.49799 |
|  | Zn66_ppm | 49.07467499 | 24.67689718 | 36.87579 |
| Wild type 4 | Fe56_ppm | 30.55253144 | 28.13670101 | 29.34462 |
|  | Cu63_ppm | 10.92706818 | 11.44964021 | 11.18835 |
|  | Zn66_ppm | 27.39776815 | 27.6259467 | 27.51186 |
| Atp7a<sup>ΔDbh</sup>_1 | Fe56_ppm | 17.03560611 | 16.06287638 | 16.54924 |
|  | Cu63_ppm | 4.314214862 | 3.69060261 | 4.002409 |
|  | Zn66_ppm | 11.35299273 | 10.63358884 | 10.99329 |
| Atp7a<sup>ΔDbh</sup>_2 | Fe56_ppm | 7.887174834 | 9.943505808 | 8.91534 |
|  | Cu63_ppm | 2.48685707 | 3.316787145 | 2.901822 |
|  | Zn66_ppm | 7.346034496 | 8.684080172 | 8.015057 |
| Atp7a<sup>ΔDbh</sup>_3 | Fe56_ppm | 15.37261191 | 15.99855668 | 15.68558 |
|  | Cu63_ppm | 5.032425899 | 4.320818919 | 4.676622 |
|  | Zn66_ppm | 12.05653024 | 15.84153393 | 13.94903 |
| Atp7b<sup>ΔDbh</sup>_1 | Fe56_ppm | 41.54731144 | 34.726308 | 38.13681 |
|  | Cu63_ppm | 14.85223744 | 8.853790876 | 11.85301 |
|  | Zn66_ppm | 38.09642812 | 32.34389281 | 35.22016 |
| Atp7b<sup>ΔDbh</sup>_2 | Fe56_ppm | 33.00461541 | 34.67047944 | 33.83755 |
|  | Cu63_ppm | 13.94175298 | 17.03608233 | 15.48892 |
|  | Zn66_ppm | 38.49497998 | 39.02257119 | 38.75878 |
| Atp7b<sup>ΔDbh</sup>_3 | Fe56_ppm | 42.87421338 | 38.52041995 | 40.69732 |
|  | Cu63_ppm | 12.99967127 | 8.379113077 | 10.68939 |
|  | Zn66_ppm | 34.38534161 | 34.52487674 | 34.45511 |

**Table S3:**
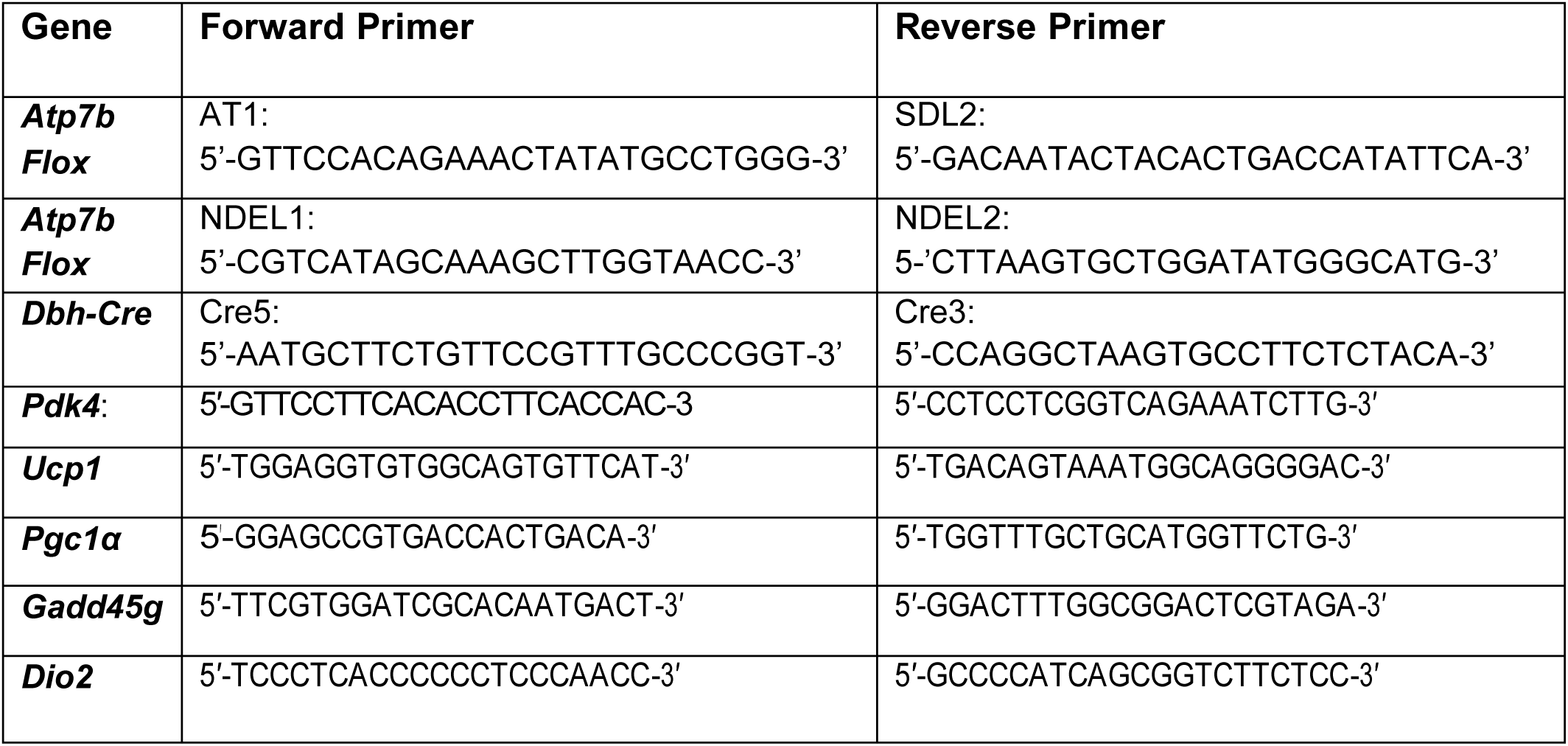
Primers used for mouse genotyping and qPCR.

**Table S4.**
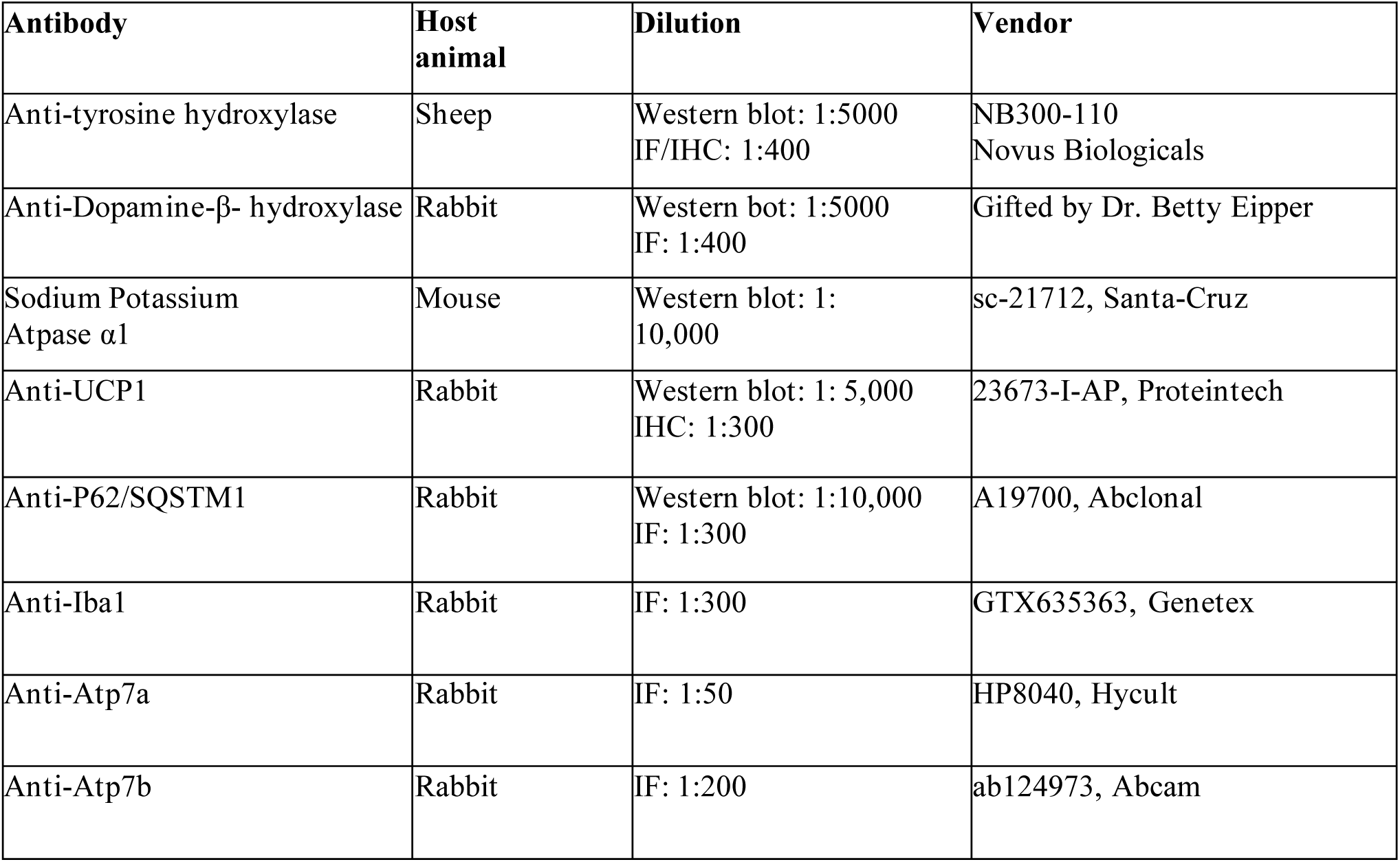
Antibodies used in immunofluorescent and immunoblot experiments.

## References

Arnold, A.C., E.M. Garland, J.E. Celedonio, S.R. Raj, N.N. Abumrad, I. Biaggioni, D. Robertson, J.M. Luther, and C.A. Shibao. 2017. Hyperinsulinemia and Insulin Resistance in Dopamine beta-Hydroxylase Deficiency. J Clin Endocrinol Metab 102:10–14.

Burman, J.L., L. Bourbonniere, J. Philie, T. Stroh, S.Y. Dejgaard, J.F. Presley, and P.S. McPherson. 2008. Scyl1, mutated in a recessive form of spinocerebellar neurodegeneration, regulates COPI-mediated retrograde traffic. J Biol Chem 283:22774–22786.

Cater, J.H., J.R. Kumita, R. Zeineddine Abdallah, G. Zhao, A. Bernardo-Gancedo, A. Henry, W. Winata, M. Chi, B.S.F. Grenyer, M.L. Townsend, M. Ranson, C.S. Buhimschi, D.S. Charnock-Jones, C.M. Dobson, M.R. Wilson, I.A. Buhimschi, and A.R. Wyatt. 2019. Human pregnancy zone protein stabilizes misfolded proteins including preeclampsia- and Alzheimer’s-associated amyloid beta peptide. Proc Natl Acad Sci U S A 116:6101–6110.

Czlonkowska, A., T. Litwin, P. Dusek, P. Ferenci, S. Lutsenko, V. Medici, J.K. Rybakowski, K.H. Weiss, and M.L. Schilsky. 2018. Wilson disease. Nat Rev Dis Primers 4:21.

Dusek, P., T. Litwin, and A. Czlonkowska. 2019. Neurologic impairment in Wilson disease. Ann Transl Med 7:S64.

Finlin, B.S., H. Memetimin, A.L. Confides, I. Kasza, B. Zhu, H.J. Vekaria, B. Harfmann, K.A. Jones, Z.R. Johnson, P.M. Westgate, C.M. Alexander, P.G. Sullivan, E.E. Dupont-Versteegden, and P.A. Kern. 2018. Human adipose beiging in response to cold and mirabegron. JCI Insight 3:

Fujisawa, C., H. Kodama, Y. Sato, M. Mimaki, M. Yagi, H. Awano, M. Matsuo, H. Shintaku, S. Yoshida, M. Takayanagi, M. Kubota, A. Takahashi, and Y. Akasaka. 2022. Early clinical signs and treatment of Menkes disease. Mol Genet Metab Rep 31:100849.

Gallagher, E.R., and E.L.F. Holzbaur. 2023. The selective autophagy adaptor p62/SQSTM1 forms phase condensates regulated by HSP27 that facilitate the clearance of damaged lysosomes via lysophagy. Cell Rep 42:112037.

Garside, J.C., E.W. Livingston, J.E. Frank, H. Yuan, and R.T. Branca. 2022. In vivo imaging of brown adipose tissue vasculature reactivity during adrenergic stimulation of non-shivering thermogenesis in mice. Sci Rep 12:21383.

Huang, Z., L. Zhong, J.T.H. Lee, J. Zhang, D. Wu, L. Geng, Y. Wang, C.M. Wong, and A. Xu. 2017. The FGF21-CCL11 Axis Mediates Beiging of White Adipose Tissues by Coupling Sympathetic Nervous System to Type 2 Immunity. Cell Metab 26:493–508 e494.

Kaeser-Pebernard, S., C. Vionnet, M. Mari, D.S. Sankar, Z. Hu, C. Roubaty, E. Martinez-Martinez, H. Zhao, M. Spuch-Calvar, A. Petri-Fink, G. Rainer, F. Steinberg, F. Reggiori, and J. Dengjel. 2022. mTORC1 controls Golgi architecture and vesicle secretion by phosphorylation of SCYL1. Nat Commun 13:4685.

Karczewski, K.J., L.C. Francioli, G. Tiao, B.B. Cummings, J. Alföldi, Q. Wang, R.L. Collins, K.M. Laricchia, A. Ganna, D.P. Birnbaum, L.D. Gauthier, H. Brand, M. Solomonson, N.A. Watts, D. Rhodes, M. Singer-Berk, E.M. England, E.G. Seaby, J.A. Kosmicki, R.K. Walters, K. Tashman, Y. Farjoun, E. Banks, T. Poterba, A. Wang, C. Seed, N. Whiffin, J.X. Chong, K.E. Samocha, E. Pierce-Hoffman, Z. Zappala, A.H. O’Donnell-Luria, E.V. Minikel, B. Weisburd, M. Lek, J.S. Ware, C. Vittal, I.M. Armean, L. Bergelson, K. Cibulskis, K.M. Connolly, M. Covarrubias, S. Donnelly, S. Ferriera, S. Gabriel, J. Gentry, N. Gupta, T. Jeandet, D. Kaplan, C. Llanwarne, R. Munshi, S. Novod, N. Petrillo, D. Roazen, V. Ruano-Rubio, A. Saltzman, M. Schleicher, J. Soto, K. Tibbetts, C. Tolonen, G. Wade, M.E. Talkowski, C. Genome Aggregation Database, C.A. Aguilar Salinas, T. Ahmad, C.M. Albert, D. Ardissino, G. Atzmon, J. Barnard, L. Beaugerie, E.J. Benjamin, M. Boehnke, L.L. Bonnycastle, E.P. Bottinger, D.W. Bowden, M.J. Bown, J.C. Chambers, J.C. Chan, D. Chasman, J. Cho, M.K. Chung, B. Cohen, A. Correa, D. Dabelea, M.J. Daly, D. Darbar, R. Duggirala, J. Dupuis, P.T. Ellinor, R. Elosua, J. Erdmann, T. Esko, M. Färkkilä, J. Florez, A. Franke, G. Getz, B. Glaser, S.J. Glatt, D. Goldstein, C. Gonzalez, L. Groop, C. Haiman, C. Hanis, M. Harms, M. Hiltunen, M.M. Holi, C.M. Hultman, M. Kallela, J. Kaprio, S. Kathiresan, B.-J. Kim, Y.J. Kim, G. Kirov, J. Kooner, S. Koskinen, H.M. Krumholz, S. Kugathasan, S.H. Kwak, M. Laakso, T. Lehtimäki, R.J.F. Loos, S.A. Lubitz, R.C.W. Ma, D.G. MacArthur, J. Marrugat, K.M. Mattila, S. McCarroll, M.I. McCarthy, D. McGovern, R. McPherson, J.B. Meigs, O. Melander, A. Metspalu, B.M. Neale, P.M. Nilsson, M.C. O’Donovan, D. Ongur, L. Orozco, M.J. Owen, C.N.A. Palmer, A. Palotie, K.S. Park, C. Pato, A.E. Pulver, N. Rahman, A.M. Remes, J.D. Rioux, S. Ripatti, D.M. Roden, D. Saleheen, V. Salomaa, N.J. Samani, J. Scharf, H. Schunkert, M.B. Shoemaker, P. Sklar, H. Soininen, H. Sokol, T. Spector, P.F. Sullivan, J. Suvisaari, E.S. Tai, Y.Y. Teo, T. Tiinamaija, M. Tsuang, D. Turner, T. Tusie-Luna, E. Vartiainen, M.P. Vawter, J.S. Ware, H. Watkins, R.K. Weersma, M. Wessman, J.G. Wilson, R.J. Xavier, B.M. Neale, M.J. Daly, and D.G. MacArthur. 2020. The mutational constraint spectrum quantified from variation in 141,456 humans. Nature 581:434–443.

Kuleshov, M.V., M.R. Jones, A.D. Rouillard, N.F. Fernandez, Q. Duan, Z. Wang, S. Koplev, S.L. Jenkins, K.M. Jagodnik, A. Lachmann, M.G. McDermott, C.D. Monteiro, G.W. Gundersen, and A. Ma’ayan. 2016. Enrichr: a comprehensive gene set enrichment analysis web server 2016 update. Nucleic Acids Res 44:W90–97.

Lee, Y.H., A.P. Petkova, E.P. Mottillo, and J.G. Granneman. 2012. In vivo identification of bipotential adipocyte progenitors recruited by beta3-adrenoceptor activation and high-fat feeding. Cell Metab 15:480–491.

Litwin, T., A. Antos, J. Bembenek, and A. Cz Onkowska. 2024. Neurological Deterioration in Wilson’s Disease-Types, Etiology, Course, and Management. Discov Med 36:646–654.

Livak, K.J., and T.D. Schmittgen. 2001. Analysis of relative gene expression data using real-time quantitative PCR and the 2(-Delta Delta C(T)) Method. Methods 25:402–408.

Lutsenko, S., S. Roy, and P. Tsvetkov. 2024. Mammalian Copper Homeostasis: Physiologic Roles and Molecular Mechanisms. Physiol Rev

Lutsenko, S., C. Washington-Hughes, M. Ralle, and K. Schmidt. 2019. Copper and the brain noradrenergic system. J Biol Inorg Chem 24:1179–1188.

Mohr, I., and K.H. Weiss. 2019. Current anti-copper therapies in management of Wilson disease. Ann Transl Med 7:S69.

Muchenditsi, A., H. Yang, J.P. Hamilton, L. Koganti, F. Housseau, L. Aronov, H. Fan, H. Pierson, A. Bhattacharjee, R. Murphy, C. Sears, J. Potter, C.R. Wooton-Kee, and S. Lutsenko. 2017. Targeted inactivation of copper transporter Atp7b in hepatocytes causes liver steatosis and obesity in mice. Am J Physiol Gastrointest Liver Physiol 313:G39–G49.

Pang, Z., G. Zhou, J. Ewald, L. Chang, O. Hacariz, N. Basu, and J. Xia. 2022. Using MetaboAnalyst 5.0 for LC-HRMS spectra processing, multi-omics integration and covariate adjustment of global metabolomics data. Nat Protoc 17:1735–1761.

Park, N.Y., D.S. Jo, J.Y. Yang, J.E. Bae, J.B. Kim, Y.H. Kim, S.H. Kim, P. Kim, D.S. Lee, T. Yoshimori, E.K. Jo, E. Yeom, and D.H. Cho. 2025. Activation of lysophagy by a TBK1-SCF(FBXO3)-TMEM192-TAX1BP1 axis in response to lysosomal damage. Nat Commun 16:1109.

Ramani, P.K., and B. Parayil Sankaran. 2024. Menkes Disease. In StatPearls. Treasure Island (FL).

Sarkar, B., K. Lingertat-Walsh, and J.T. Clarke. 1993. Copper-histidine therapy for Menkes disease. J Pediatr 123:828–830.

Sarver, D.C., A.N. Stewart, S. Rodriguez, H.C. Little, S. Aja, and G.W. Wong. 2020. Loss of CTRP4 alters adiposity and food intake behaviors in obese mice. Am J Physiol Endocrinol Metab 319:E1084–E1100.

Sayar-Atasoy, N., C. Laule, I. Aklan, H. Kim, Y. Yavuz, T. Ates, I. Coban, F. Koksalar-Alkan, J. Rysted, D. Davis, U. Singh, M.I. Alp, B. Yilmaz, H. Cui, and D. Atasoy. 2023. Adrenergic modulation of melanocortin pathway by hunger signals. Nat Commun 14:6602.

Schmidt, K., M. Ralle, T. Schaffer, S. Jayakanthan, B. Bari, A. Muchenditsi, and S. Lutsenko. 2018. ATP7A and ATP7B copper transporters have distinct functions in the regulation of neuronal dopamine-beta-hydroxylase. J Biol Chem 293:20085–20098.

Schwarz, L.A., and L. Luo. 2015. Organization of the locus coeruleus-norepinephrine system. Curr Biol 25:R1051–R1056.

Sciolino, N.R., M. Hsiang, C.M. Mazzone, L.R. Wilson, N.W. Plummer, J. Amin, K.G. Smith, C.A. McGee, S.A. Fry, C.X. Yang, J.M. Powell, M.R. Bruchas, A.V. Kravitz, J.D. Cushman, M.J. Krashes, G. Cui, and P. Jensen. 2022. Natural locus coeruleus dynamics during feeding. Sci Adv 8:eabn9134.

Smythe, G.A., H.S. Grunstein, J.E. Bradshaw, M.V. Nicholson, and P.J. Compton. 1984. Relationships between brain noradrenergic activity and blood glucose. Nature 308:65–67.

Tominaga, T., M. Goto, T. Onoue, A. Mizoguchi, M. Sugiyama, T. Tsunekawa, D. Hagiwara, Y. Morishita, Y. Ito, S. Iwama, H. Suga, R. Banno, and H. Arima. 2017. Sequestosome 1 (SQSTM1/p62) maintains protein folding capacity under endoplasmic reticulum stress in mouse hypothalamic organotypic culture. Neurosci Lett 656:103–107.

Tumer, Z., and L.B. Moller. 2010. Menkes disease. Eur J Hum Genet 18:511–518.

Tusher, V.G., R. Tibshirani, and G. Chu. 2001. Significance analysis of microarrays applied to the ionizing radiation response. Proc Natl Acad Sci U S A 98:5116–5121.

Tyanova, S., T. Temu, P. Sinitcyn, A. Carlson, M.Y. Hein, T. Geiger, M. Mann, and J. Cox. 2016. The Perseus computational platform for comprehensive analysis of (prote)omics data. Nat Methods 13:731–740.

Walters, J.M., G.M. Ward, J. Barton, R. Arackal, R.C. Boston, J.D. Best, and F.P. Alford. 1997. The effect of norepinephrine on insulin secretion and glucose effectiveness in non-insulin-dependent diabetes. Metabolism 46:1448–1453.

Washington-Hughes, C.L., S. Roy, H.K. Seneviratne, S.S. Karuppagounder, Y. Morel, J.W. Jones, A. Zak, T. Xiao, T.N. Boronina, R.N. Cole, N.N. Bumpus, C.J. Chang, T.M. Dawson, and S. Lutsenko. 2023. Atp7b-dependent choroid plexus dysfunction causes transient copper deficit and metabolic changes in the developing mouse brain. PLoS Genet 19:e1010558.

Xiao, T., C.M. Ackerman, E.C. Carroll, S. Jia, A. Hoagland, J. Chan, B. Thai, C.S. Liu, E.Y. Isacoff, and C.J. Chang. 2018. Copper regulates rest-activity cycles through the locus coeruleus-norepinephrine system. Nat Chem Biol 14:655–663.

Yamaguchi, H., F.W. Hopf, S.B. Li, and L. de Lecea. 2018. In vivo cell type-specific CRISPR knockdown of dopamine beta hydroxylase reduces locus coeruleus evoked wakefulness. Nat Commun 9:5211.

